# Proteomic identification and structural basis for the interaction between sorting nexin SNX17 and PDLIM family proteins

**DOI:** 10.1101/2021.07.29.454387

**Authors:** Michael D. Healy, Joanna Sacharz, Kerrie E. McNally, Calum McConville, Vikas A. Tillu, Ryan J. Hall, Molly Chilton, Peter J. Cullen, Mehdi Mobli, Rajesh Ghai, David A. Stroud, Brett M. Collins

**Affiliations:** The University of Queensland, Institute for Molecular Bioscience, Queensland, 4072, Australia; Department of Biochemistry and Pharmacology, The Bio21 Molecular Science and Biotechnology Institute, The University of Melbourne, Parkville, Victoria, 3052, Australia; School of Biochemistry, Biomedical Sciences Building, University of Bristol, Bristol BS8 1TD, UK; MRC Laboratory of Molecular Biology, Cambridge, CB2 0QH, UK; Centre for Advanced Imaging and School of Chemistry and Molecular Biology, The University of Queensland, St. Lucia, Queensland, 4072, Australia; CSL Limited, Parkville, Victoria, 3052, Australia

**Keywords:** Commander, endosome, PDLIM, PDZ domain, Retriever, SNX17

## Abstract

The sorting nexin SNX17 controls endosome-to-cell surface recycling of diverse transmembrane cargo proteins including integrins, the amyloid precursor protein and lipoprotein receptors. This requires association with the multi-subunit Commander trafficking complex, which depends on the C-terminus of SNX17 through unknown mechanisms. Using affinity enrichment proteomics, we find that a C-terminal peptide of SNX17 is not only sufficient for Commander interaction but also associates with members of the actin-associated PDZ and LIM domain (PDLIM) family. We show that SNX17 contains a type III PSD95/Dlg/Zo1 (PDZ) binding motif (PDZbm) that binds specifically to the PDZ domains of PDLIM family proteins but not to other PDZ domains tested. The structure of the PDLIM7 PDZ domain bound to the SNX17 C-terminus was determined by NMR spectroscopy and reveals an unconventional perpendicular peptide interaction. Mutagenesis confirms the interaction is mediated by specific electrostatic contacts and a uniquely conserved proline-containing loop sequence in the PDLIM protein family. Our results define the mechanism of SNX17-PDLIM interaction and suggest that the PDLIM proteins may play a role in regulating the activity of SNX17 in conjunction with Commander and actin-rich endosomal trafficking domains.

## Introduction

Endosomes are central organelles for the trafficking and homeostasis of transmembrane proteins in the secretory and endocytic system. Protein cargos are either sorted into late endosomes and lysosomes for degradation or recycled from endosomes to various destinations such as the plasma membrane or the trans-Golgi network (TGN) within tubulovesicular transport carriers. These are formed by several different protein machineries including sorting nexins (SNXs) with Bin/Amphiphysin/Rvs (BAR) domains and the Retromer complex. Retromer consists of three proteins, VPS35–VPS26–VPS29, and associates with transmembrane cargo molecules via adaptor proteins from the SNX family including SNX3 and SNX27 (Chen et al., 2019; Weeratunga et al., 2020).

A large number of receptors and adhesion proteins such as lipoprotein receptor family members LDLR and LRP1 (Burden et al., 2004; Farfan et al., 2013; van Kerkhof et al., 2005), the amyloid precursor protein (APP) (Lee et al., 2008), P-selectin and the α5β1 integrin complex (Bottcher et al., 2012; Steinberg et al., 2012; Tseng et al., 2014) are recycled from endosomes to the plasma membrane via interaction with the ubiquitous SNX family member SNX17, or its paralogue SNX31 in bladder urothelial cells (McNally et al., 2017; Tseng et al., 2014; Vieira et al., 2014). This occurs via binding of the SNX17 4.1/ezrin/radixin/moesin (FERM) domain to ΦxNxx[YF] motifs (where Φ is a hydrophobic amino acid) within the cytoplasmic domains of the receptors (Ghai et al., 2013; Ghai et al., 2011). Recent work showed that SNX17 and SNX31 act in concert with a large multi-subunit assembly called Commander (McNally et al., 2017), which is composed of two complexes named Retriever and CCC (Chen et al., 2019; Mallam and Marcotte, 2017; McNally and Cullen, 2018) (see **Fig. 6C** for a cartoon summary of key interactions). Retriever is predicted to be structurally similar to Retromer and is composed of the three subunits VPS35L/C16orf62, VPS26C/DSCR3 and VPS29 (McNally and Cullen, 2018). The CCC complex is comprised of ten members of the Copper Metabolism MURR1 domain (COMMD) family Commd1-10, and two coiled-coil domain-containing proteins CCDC93 and CCDC22. How SNX17 and Commander mediate endosomal recycling is unknown, but it is facilitated by interactions with the Arp2/3 activating Wiskott–Aldrich syndrome protein and SCAR homologue (WASH) complex (Bartuzi et al., 2016; Campion et al., 2018; McNally et al., 2017; Phillips-Krawczak et al., 2015; Simonetti and Cullen, 2019; Singla et al., 2019). Dynamic actin polymerisation at the endosome appears to be essential, and depletion of Commander proteins strongly perturbs these dynamics leading to endosomal accumulation of WASH, actin, and transmembrane cargos (Campion et al., 2018; Singla et al., 2019).

The interaction between SNX17 and Commander is disrupted by a single point mutation of the SNX17 C-terminal Leu (L470G) (McNally et al., 2017), and in this work we set out to study the mechanism by which SNX17 engages with Commander to mediate endosomal cargo recycling. Starting with affinity enrichment mass-spectrometry we found a short peptide derived from the C-terminus of SNX17 is sufficient to recruit the entire Commander assembly composed of Retriever and CCC (McNally et al., 2017). Notably however we also identified specific interactions with several members of the PDZ and LIM domain-containing (PDLIM) protein family; including PDLIM1, PDLIM4, and PDLIM7. PDLIM proteins are multi-domain scaffolds that are typically associated with actin and its accessory proteins such as α-actinins and myotilin (Bauer et al., 2000; Huang et al., 2020; Katzemich et al., 2013; Klaavuniemi et al., 2009; Klaavuniemi et al., 2004; Krcmery et al., 2010; Liu et al., 2015; Tamura et al., 2007; te Velthuis and Bagowski, 2007; Vallenius et al., 2000; Vallenius et al., 2004; von Nandelstadh et al., 2009; Zheng et al., 2010). We show that the purified PDZ domains of the PDLIM family members bind specifically *in vitro* to the SNX17 C-terminal peptide, and that SNX17 can bind PDLIM7 in cells. The C-terminal sequence of SNX17 (DEDL) conforms to a Type III PDZ binding motif (PDZbm) and the solution structure of the PDLIM7 PDZ domain bound to this sequence reveals an unconventional mode of peptide interaction. Canonically, PDZbm peptides form an extended β-strand between the second β-strand (βB) and second α-helix (αB) of the PDZ domain, with their C-terminal carboxyl group forming hydrogen bonds with the main chain amide groups of the PDZ domain loop preceding the βB strand (Lee and Zheng, 2010; Subbaiah et al., 2011; Ye and Zhang, 2013). The carboxylate-binding loop typically has the sequence G<λG<λx (where x is any amino acid residue and Φ is a hydrophobic residue). PDLIM proteins instead possess a unique loop sequence, PWGFR, that we find is essential for SNX17 PDZbm binding in this atypical upright conformation. Altogether this work identifies a novel association between the PDLIM family proteins and the SNX17 endosomal adaptor proteins and suggests a potential role in actin and Commander-dependent trafficking and signalling.

## Results

### The C-terminus of SNX17 contains a Type III PDZ binding motif that interacts with PDLIM family proteins

Initially we hoped to gain insights into the mechanism by which SNX17 interacts with Commander by performing an unbiased proteomic analysis, comparing the proteomes of a biotinylated synthetic peptide derived from the C-terminus of SNX17 (^457^HGNFAFEGIGDEDL^470^) with a mutant peptide sequence (^457^HGNFAFEGIGDED<u>G</u>^470^). The L470G mutation in SNX17 has previously been shown to block the association with Commander subunits following immunoprecipitation (McNally et al., 2017). Using these peptides as baits in an affinity enrichment mass-spectrometry experiment from HEK293T cell lysates, all known components of both the Retriever and CCC sub-complexes (except for Commd6, which was not quantified at background levels, suggesting a detection issue rather than a lack of enrichment) were specifically enriched by the native sequence relative to the mutant, indicating that the C-terminal tail of SNX17 is both required and sufficient for Commander association (**Fig. 1A; Table S1**).

**Figure 1.**
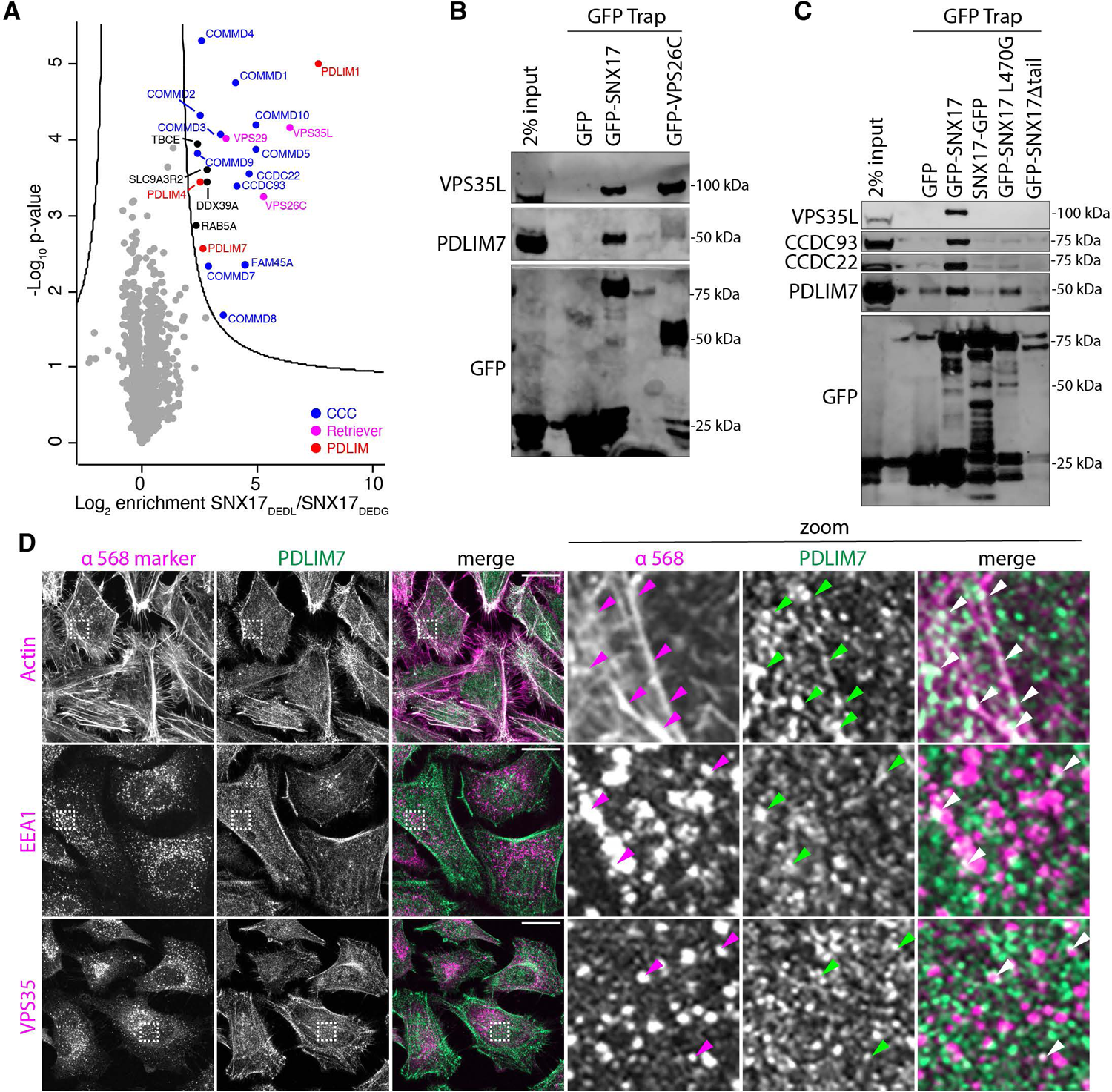
Identification of SNX17 C-terminal interacting proteins. **(A)** Volcano plot of identified interactors enriched in pull-downs with biotinylated SNX17 C-terminal peptide (HGNFAFEGIGDEDL) versus a control mutant SNX17 peptide (HGNFAFEGIGDEDG). The entire Commander assembly is identified (CCC subunits in blue and Retriever subunits in magenta) as are several members of the PDLIM family (red). **(B)** GFP traps of GFP-tagged SNX17 and VPS26C from HEK293 cells. Bound VPS35L and PDLIM7 were detected by Western blot with specific antibodies. Both SNX17 and VPS26C bind endogenous Commander subunit VPS35L, while PDLIM7 is specifically associated with SNX17. **(C)** GFP traps of SNX17 wild-type protein and C-terminal mutants with either N- or C-terminal GFP tags from HEK293 cells. Bound proteins were detected by Western blot. Either mutation or tagging at the C-terminus disrupts SNX17 binding to the Commander complex and the PDLIM7 protein. **(D)** The colocalisation of endogenous PDLIM7 (green) with actin or early endosome markers VPS35 and EEA1 (magenta) was assessed in HeLa cells by immunofluorescence confocal microscopy. Arrows indicate instances of PDLIM7 puncta overlapping with endosomal compartments. Scale bars represent 10 µm. White dashed boxes represent areas that have been zoomed in.

The C-terminal sequence of SNX17 (DEDL) conforms to the consensus of a Type III PDZ binding motif ([ED]×Φ); where Φ is a hydrophobic side chain at the C-terminus) and is highly conserved throughout evolution, as well as being conserved in the bladder-specific paralogue SNX31 (see **Fig. 6B**). Although Commander lacks any known PDZ domain containing subunits, the perturbed association following the single L470G substitution at the C-terminus of SNX17 strongly suggests a role for a PDZ domain-containing scaffold protein(s) in complex formation. Strikingly, in addition to Commander a select number of novel interacting proteins were also identified in SNX17_DEDL_ eluates, including three proteins PDLIM1, PDLIM4 and PDLIM7 that are members of the PDZ and LIM domain-containing family of PDLIM proteins (**Fig. 1A; Table. S1**). We confirmed the interaction in HEK293 cells expressing GFP-SNX17 by co-immunoprecipitation, with western blots of proteins isolated by GFP-Trap showing GFP-SNX17 association with both the endogenous VPS35L subunit of Retriever and endogenous PDLIM7 (**Fig. 1B**). In contrast, while GFP-VPS26C interacts with VPS35L as expected, it does not appear to associate with PDLIM7 under these conditions. In line with the importance of the C-terminal Leu470 residue, when the GFP-tag is moved to the C-terminus of SNX17 the binding to both Commander and PDLIM7 is reduced (**Fig. 1C**).

We next examined the localisation of PDLIM7 in HeLa cells by immunofluorescence microscopy (**Fig. 1D**). As observed previously (Bauer et al., 2000; D’Cruz et al., 2016; Elbediwy et al., 2018; Klein et al., 2018; Shi et al., 2020; Tamura et al., 2007; Urban et al., 2016; Xia et al., 1997; Yi et al., 2014) we see significant colocalization with the actin cytoskeleton along with other cytoplasmic punctate structures (**Fig. 1D**). Using both EEA1 and VPS35 as markers we see a small degree of colocalisation between PDLIM7 and the early endosomal compartments, with a minor population of overlapping structures observed. Similar results were seen following transient expression of GFP-tagged PDLIM7 in HeLa cells (**Fig. S1A**), or when analysing the localisation of endogenous PDLIM7 in RPE1 cells (**Fig. S1B and S1C**). In addition, we see a similar localisation phenotype for PDLIM1-GFP (**Fig. S1D**). This suggests that association of PDLIM7 with SNX17 is transient and/or only occurring on a sub-population of endosomes, and likely reflects a specialised function of PDLIM proteins alongside their major roles in actin cytoskeleton regulation.

### The SNX17 peptide binds specifically to the PDZ domain of PDLIM proteins *in vitro*

The human PDLIM protein family has seven members, PDLIM1 to PDLIM7 (**Fig. 2A**). These have a number of important scaffolding functions for proteins including transcription factors (Camarata et al., 2006; Krause et al., 2004) and pathogenic effectors (Dhanda et al., 2021; Fu et al., 2010; Yan et al., 2009; Yi et al., 2014), with a prominent role in organisation of the actin cytoskeleton by interacting directly with actin and α-actinin in the Z-discs of muscle cells and actin stress fibres (Bauer et al., 2000; Katzemich et al., 2013; Klaavuniemi et al., 2009; Klaavuniemi et al., 2004; Liu et al., 2015; Tamura et al., 2007; te Velthuis and Bagowski, 2007; Vallenius et al., 2000; Vallenius et al., 2004; von Nandelstadh et al., 2009; Zheng et al., 2010). They all possess an N-terminal PDZ domain followed by a long unstructured sequence connected to C-terminal LIM (Lin-11, Isl1 and Mec-3) domains, and can be divided into two sub-families; the ALP (Actinin-associated LIM protein) homologues with a single LIM domain, and the Enigma homologues with three tandem LIM domains, (**Fig. 2A; Fig. S2; Fig. S3**).

**Figure 2.**
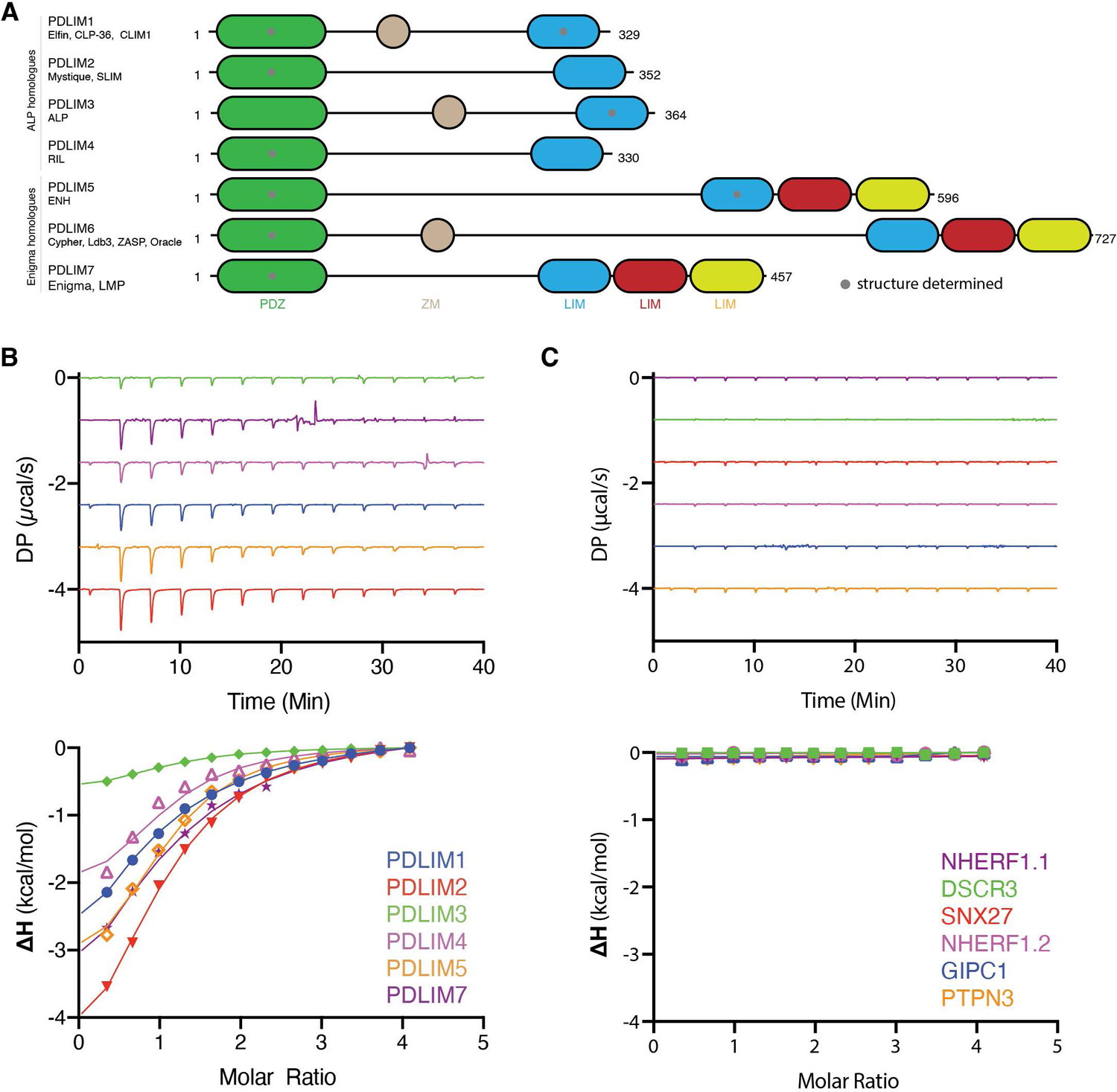
Direct interaction of the C-terminal SNX17 PDZbm with PDLIM family proteins. **(A)** Domain architecture of the PDLIM family proteins drawn to scale. The ALP homologues PDLIM1, PDLIM2, PDLIM3/ALP and PDLIM4 possess a single LIM domain at their C-terminus, while the Enigma homologues PDLIM5, PDLIM6 and PDLIM7/Enigma have three tandem LIM domains. Grey dots indicate domains for which crystal or NMR structures have been determined. Sequence alignments of the PDLIM proteins are provided in **Fig. S2** and the sites of identified post-translational modifications of PDLIM proteins are shown in **Fig. S3. (B)** Binding of PDZ domains from PDLIM family members to the SNX17 peptide measured by ITC. The top panels show raw data while the bottom shows the normalised and integrated binding curves. **(C)** No binding of the SNX17 C-terminal peptide was observed to a variety of other PDZ domains tested.

To confirm the direct interaction between SNX17 and the PDLIM proteins we purified the PDZ domains of six of the seven human PDLIM proteins and tested their binding to the SNX17 peptide using isothermal titration calorimetry (ITC). SNX17 interacts directly with all six PDLIM family members with modest affinities (*K*_d_s) of between 10 to 40 μM. (**Fig. 2B; Table 1**). The selective isolation of PDLIM1, PDLIM4 and PDLIM7 in affinity enrichment proteomics likely reflects differential expression levels of the various family members rather than different affinities; for example PDLIM3 and PDLIM6 are almost exclusively expressed in heart and skeletal muscle tissues (Faulkner et al., 1999; Passier et al., 2000; Pomies et al., 1999; Wu et al., 2016; Xia et al., 1997; Zhou et al., 1999). In contrast to PDLIM binding, the PDZ domains from control proteins including GIPC1, NHERF1 (domains 1 and 2), PTPN3, and SNX27 showed no detectable interaction with the SNX17 peptide under these conditions (**Fig. 2C; Table 1**). In addition we did not observe significant binding *in vitro* of the SNX17 peptide to the Retriever subunit VPS26C, which has previously been speculated to be involved in the interaction because its depletion perturbs SNX17 association with Retriever (McNally et al., 2017). Taken together these results show that the C-terminal Type III PDZbm in SNX17 can selectively bind the PDLIM family of proteins in cells and *in vitro*.

**Table 1.** Thermodynamic binding parameters of PDZ domains with PDZbm peptides^a^.

| Protein (cell) | Peptide (syringe) | $K_d$ (μM) | $\Delta H$ (kcal/mol) | $\Delta G$ (kcal/mol) | $-T\Delta S$ (kcal/mol) |
| --- | --- | --- | --- | --- | --- |
| PDLIM1 | SNX17 | 36.0 ± 11.8 | -4.02 ± 1.71 | -6.40 ± 0.262 | -2.38 ± 1.97 |
| PDLIM2 | SNX17 | 23.4 ± 0.12 | -5.57 ± 0.28 | -6.57 ± 0.003 | -0.83 ± 0.28 |
| PDLIM3 | SNX17 | 19.0 ± 1.80 | -0.91 ± 0.13 | -6.70 ± 0.064 | -5.80 ± 0.16 |
| PDLIM4 | SNX17 | 11.2 ± 3.07 | -2.31 ± 0.13 | -7.07 ± 0.165 | -4.76 ± 0.30 |
| PDLIM5 | SNX17 | 25.3 ± 5.94 | -4.76 ± 0.77 | -6.56 ± 0.141 | -1.79 ± 0.91 |
| PDLIM7 | SNX17 | 18.2 ± 1.82 | -4.27 ± 0.60 | -6.74 ± 0.060 | -2.46 ± 0.66 |
| PDLIM7(R17D) | SNX17 | NB |  |  |  |
| PDLIM7(H63A) | SNX17 | 16.6 ± 1.19 | -3.62 ± 0.40 | -6.79 ± 0.046 | -3.17 ± 0.44 |
| PDLIM7(Q67A) | SNX17 | 13.3 ± 3.08 | -5.54 ± 0.49 | -6.96 ± 0.158 | -1.42 ± 0.83 |
| PDLIM7(P11E) | SNX17 | 32.7 ± 4.40 | -3.01 ± 0.70 | -6.38 ± 0.085 | -3.37 ± 0.78 |
| PDLIM7(P13G) | SNX17 | NB |  |  |  |
| PDLIM7(P11E/P13G) | SNX17 | NB |  |  |  |
| PDLIM7 | SNX17(L470G) | NB |  |  |  |
| PDLIM7 | SNX17(D467S/E468S/D469S) | NB |  |  |  |
| PDLIM7 | SNX17(E468R) | 50.8 ± 5.2 | -4.38 ± 0.47 | -6.10 ± 0.068 | -1.72 ± 0.53 |
| NHERF1.1 | SNX17 | NB |  |  |  |
| NHERF1.2 | SNX17 | NB |  |  |  |
| SNX27 | SNX17 | NB |  |  |  |
| GIPC1 | SNX17 | NB |  |  |  |
| PTPN3 | SNX17 | NB |  |  |  |
| DSCR3 | SNX17 | NB |  |  |  |
| PDLIM7 | α-actinin-1 | 62.8 ± 11.10 | -5.80 ± 0.79 | -5.98 ± 0.106 | -0.19 ± 0.90 |
| PDLIM7(R17D) | α-actinin-1 | NB | | | |
| PDLIM7(H63A) | α-actinin-1 | 106.0 ± 16.8 | -5.06 ± 0.50 | -5.65 ± 0.010 | -0.60 ± 0.60 |
| PDLIM7 | Myotilin | 7.3 ± 0.44 | -8.16 ± 0.28 | -7.30 ± 0.004 | 0.86 ± 0.29 |
| PDLIM7(R17D) | Myotilin | NB |  |  |  |
| PDLIM7(H63A) | Myotilin | 3.7 ± 0.31 | -7.1 ± 0.18 | -7.72 ± 0.005 | -0.63 ± 0.23 |

a. Errors show SD from at least three independent experiments.

### Molecular basis of SNX17 interaction with the PDLIM7 PDZ domain

To define the mechanism of interaction between SNX17 and the PDLIM PDZ domains we used NMR spectroscopy to determine the solution structure of the PDLIM7 PDZ domain fused to the SNX17 C-terminal sequence with a long flexible linker. By fusing the SNX17 sequence to the PDLIM7 PDZ domain we could perform a full spectral assignment of the complex using ^15^N/^13^C/^1^H heteronuclear NMR. The NMR spectra were of excellent quality and allowed for a full structure determination of the complex (**Figs. 3A and 3B; Table 2; Fig. S4A**).

**Figure 3.**
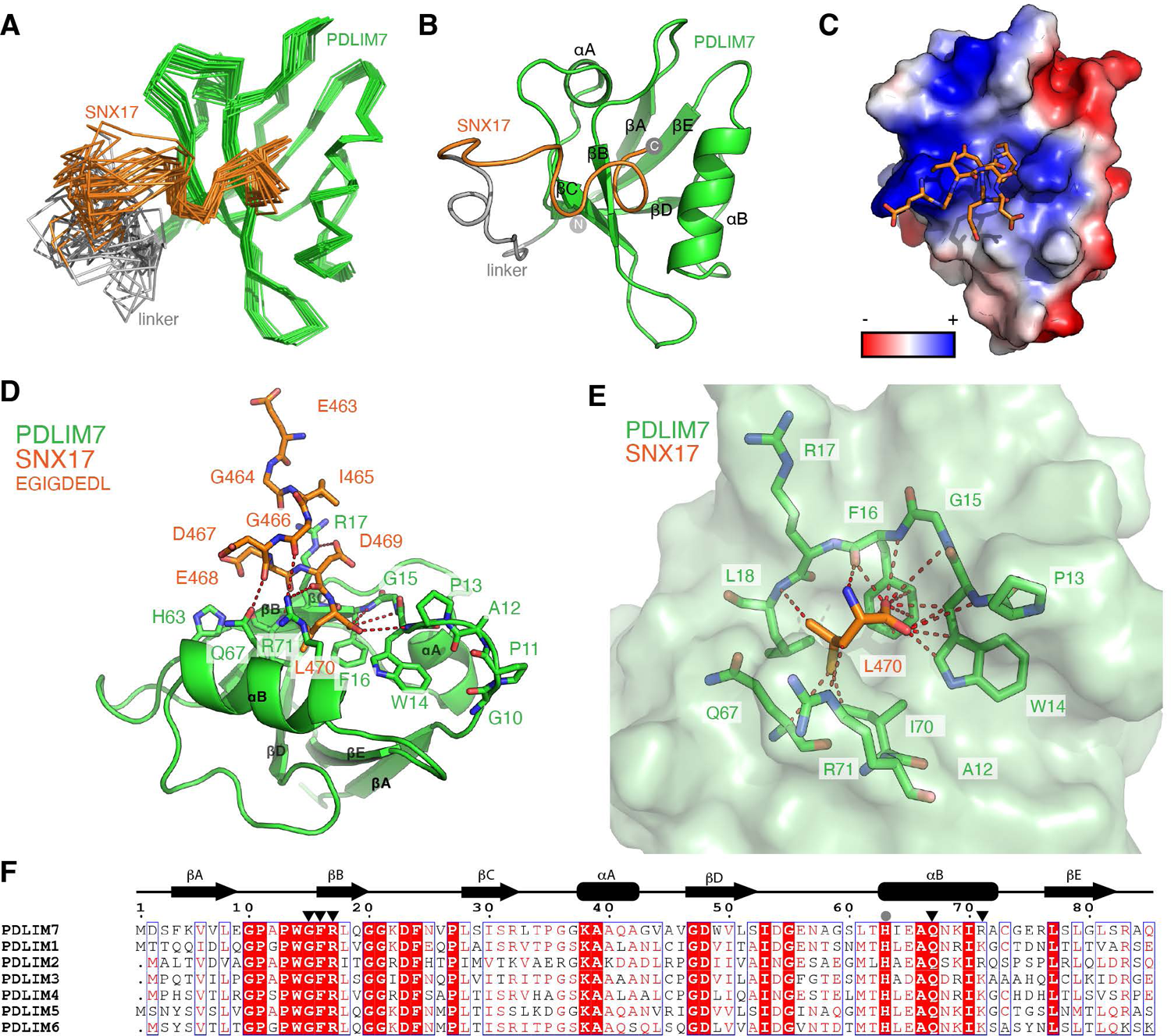
Molecular basis of SNX17 interaction with the PDLIM7 PDZ domain. **(A)** The structure of the PDLIM7 PDZ domain fused to the SNX17 N-terminal sequence was determined by NMR spectroscopy. The image shows an overall alignment of the Cα positions from the 20 lowest energy conformers, with PDLIM7 shown in green, SNX17 shown in orange, and the disordered linker sequence shown in grey. **(B)** Ribbon diagram of the PDLIM7-SNX17 complex with PDLIM7 secondary structure elements labelled. **(C)** Electrostatic surface potential of the PDLIM7 PDZ domain is coloured red (negatively charged) to blue (positively charged). The acidic SNX17 sequence binds to a complementary basic surface on PDLIM7. **(D)** Detailed view of the SNX17 peptide sequence (orange) and critical contacts with PDLIM7 (green). The SNX17 peptide is bound in an unconventional perpendicular orientation with respect to the PDLIM7 structure. **(E)** Close-up of the C-terminal Leu470 residue of SNX17 highlighting the extensive hydrogen bond and van der Waals interactions made with the PDLIM7 protein. (**F**) Sequence alignment of the PDZ domains of human PDLIM family members. The secondary structure of PDLIM7 is indicated above for reference, and black triangles indicate residues making direct contact with the SNX17 PDZbm peptide.

**Table 2.** Statistics for NMR structure determination of PDLIM7 PDZ domain-SNX17 PDZbm fusion protein^1^.

| **Statistics** | **Value** |
| --- | --- |
| **NMR distance and dihedral constraints<sup>2</sup>** |  |
| Total NOEs | 3886 |
| Intra-residue | 1697 |
| Inter-residue |  |
| Sequential | 1077 |
| Medium-range | 332 |
| Long-range | 780 |
| Hydrogen bond(s) | 34 |
| Dihedral angles |  |
| Backbone ($\phi$ angle) | 171 |
| **Structural statistics** |  |
| RMSD between 20 conformers (Å) |  |
| Average pairwise RMSD (Å) |  |
| Backbone (Å) (N, Cα, C) | 0.17 |
| All heavy atoms (Å) | 0.46 |
| Ramachandran analysis<sup>3</sup> |  |
| Residues in most favoured regions (%) | 79.6 |
| Residues in additionally allowed regions (%) | 20.4 |
| Residues in generously allowed regions (%) | 0.0 |
| Residues in disallowed regions (%) | 0.0 |
| **PDB ID** | **7RM8** |
| **BMRB ID** | **30937** |

1. All statistics are given as mean ± SD.
2. Only structurally relevant restraints as defined by CYANA are included.
3. According to Molprobit (molprobit.biochem.duke.edu).

As expected from previous studies (Elkins et al., 2010), the PDZ domain of PDLIM7 (residues 1-84) forms a classical PDZ fold composed of five β-strands (βA-βE) and two α-helices (αA and αB) (**Fig. 3B**). The contacts between SNX17 and the PDLIM7 PDZ domain are exclusively mediated by the extreme C-terminal residues of SNX17 (^463^EGIGDEDL^470^), which dock into a highly conserved and positively charged surface pocket (**Fig. 3C**). The linker and upstream SNX17 sequences of the fusion protein provide a flexible extension that is long enough to allow for intramolecular peptide interaction to occur in *cis* without steric hindrance (**Figs. 3A and 3B**).

The structure of the SNX17 Type III PDZbm sequence bound to PDLIM7 shows that the C-terminal SNX17 amino acid residue Leu470 docks within the expected pocket formed by the loop preceding the βB strand (**Fig. 3D**). As seen in all known PDZ domain interactions the C-terminal carboxylate of the bound peptide is required for binding, forming direct hydrogen bonds with the backbone amides surrounding the core binding pocket (**Fig. 3E**). In PDLIM7 these main-chain amides are provided by Trp14, Gly15 and Phe16, with an additional backbone hydrogen bond formed between the amide of SNX17 Leu470 and the carboxyl group of PDLIM7 Phe16. The hydrophobic side chain of SNX17 Leu470 is nestled within a complementary pocket formed by the side chains of Phe16, Leu18 and Ile70, and the exposed aliphatic regions of Gln67 and Arg71. Furthermore, our refined structure reveals an interaction between the C-terminal hydroxyl group of SNX17 Leu470 and the aromatic ring of PDLIM7 Phe16 (**Fig. 3E; Fig. S4B**).

The vast majority of PDZ domain interactions involve a so-called ‘conventional’ binding mode, where the PDZbm peptide binds along the groove between the βB strand and αB helix, forming a β-sheet extension with the βB strand and with its C-terminal residue nestled in the core binding pocket. The C-terminal PDZbm residue is numbered ‘0’, and in the most common Type I sequences ([ST]xΦ) the side chain at position −2 is a Ser/Thr that mediates a hydrogen bond to a His side chain in the PDZ domain at the beginning of the αB helix (His63 in PDLIM7). The SNX17 Type III PDZbm however adopts an unconventional ‘upright’ conformation when bound to PDLIM7. Here the C-terminal Leu470 at position 0 is bound as expected, but the upstream residues are oriented away from the groove between strand βB and helix αB (**Fig. 3D**). A consequence of this is that there are a relatively small number of main-chain hydrogen bonds between the PDZ domain and peptide, all of which are mediated by the C-terminal SNX17 Leu470 residue. This smaller number of main-chain interactions appears to be partly compensated by electrostatic interactions of the Glu468 and Asp469 side chains of SNX17 at the −2 and −1 positions with Arg17 in the βB strand and Arg71 in the αB helix of PDLIM7. The Arg17 side chain is strictly conserved in the PDLIM family of PDZ domains pointing to its important role in binding specificity (**Fig. 3F**).

### Comparison of the PDLIM7-SNX17 structure with other PDZ domain complexes

Given the unusual binding mode of SNX17 to the PDLIM7 PDZ domain we next compared this structure to other related PDZ domain complexes (**Fig. 4**). All Type I PDZbm sequences that adopt the conventional ‘in-groove’ binding mode invariably use their Ser or Thr side chains at position −2 to form a hydrogen bond with a conserved His in the αB helix of their respective PDZ domain partner. In contrast, the unconventional ‘upright’ binding orientation of the SNX17 Type III PDZbm to PDLIM7 is likely necessary for their interaction because the longer SNX17 Glu468 sidechain at position −2 will be sterically precluded from docking in the αB-βB groove and forming a hydrogen bond with the corresponding His63 in PDLIM7. Unconventional ‘upright’ PDZ binding modes have been seen in several previous structures reported by Elkins *et al*., 2010 (Elkins et al., 2010). These include Type I motifs from β-tropomyosin and α-actinin-1 (ESDL and ITSL respectively) bound to PDLIM2, PDLIM5 and PDLIM7, and the Type I motif from mGluR1 (SSTL) bound to GRASP (Elkins et al., 2010). In addition, the structure of the tamalin PDZ domain bound to its own C-terminal sequence (ESQL) also shows an upright orientation (Sugi et al., 2007). However, the functional significance of these ‘upright’ binding modes for the β-tropomyosin, α-actinin-1, mGluR1 and tamalin peptides is still uncertain. A significant caveat with these structures is that they all used very short peptide fusion sequences. The covalently linked C-terminal peptides are thus restricted in their flexibility and invariably bind in *trans* to neighbouring molecules in the crystal lattices; and close inspection indicates that their orientations are all highly influenced by crystal packing. Further suggesting the unconventional interactions by these Type I motifs may be artefacts of crystal packing is that the α-actinin1, mGluR1 and tamalin sequences were also found to adopt the expected conventional orientation. In the case of mGluR1 and tamalin both the conventional and unconventional binding modes are seen in the *same* crystal structure. Hence, while these structures show that it is *possible* for Type I PDZbms to adopt an upright unconventional binding mode, it is likely that the conventional in-groove binding mode is more functionally relevant.

**Figure 4.**
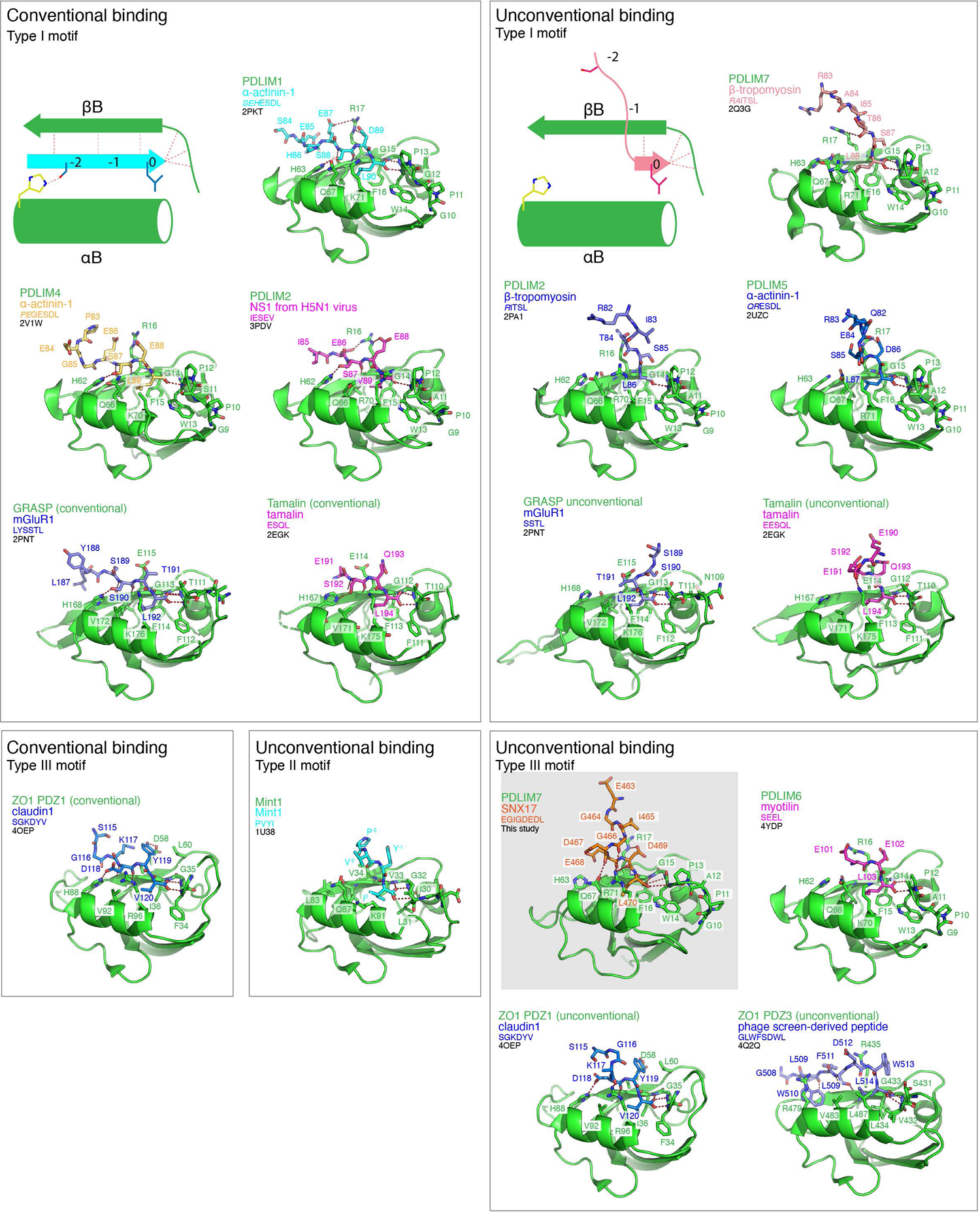
Comparison of structures of PDZ domains bound to PDZbms in both conventional and unconventional orientations. All known structures of PDLIM PDZ domains (green ribbons) in complex with PDZbms (coloured sticks) are shown. Also shown for comparison are all known examples of other Type III PDZbm structures, and of Type I PDZbm structures in an unconventional orientation. Schematic diagrams indicate how conventional binding involves formation of a β-strand in the groove between the βB strand and αB helix, while unconventional docking involves a perpendicular peptide orientation. PDLIM PDZ domains bound to various PDZbm peptides are typically in either a conventional binding conformation, parallel to the βB strand of the PDZ domain, or an unconventional upright conformation as seen for the SNX17 interaction with PDLIM7. Structures of the GRASP PDZ domain bound to a Type I motif from mGluR1 and tamalin bound to its own Type I motif show *both* conventional and unconventional orientations in the same crystal structures, where one chain of the asymmetric unit binds in the conventional manner and the second chain in the asymmetric unit binds in an unconventional manner. Therefore, it is unclear if the unconventional binding seen in these structures is an artefact of the crystal packing interactions.

In addition to the PDLIM7 structure bound to SNX17, three other structures of PDZ domains bound to Type III motifs have been determined (**Fig. 4**). The Type III motif of myotilin (SEEL) bound to PDLIM6 (Elkins et al., 2010), like SNX17 (DEDL), uses acidic sidechains in the −1 and −2 positions to form similar electrostatic interactions with the PDLIM6 Arg16 sidechain (equivalent to PDLIM7 Arg17) (Elkins et al., 2010). The structure of ZO1 PDZ domain 3 bound to an artificial phage-selected peptide (SDWL) reveals a slight variation to the unconventional binding mode (Ernst et al., 2014). ZO1 PDZ3 has an arginine, Arg479, in place of the more common His in the αB helix that interacts with Trp510 at the −5 position of the peptide. The peptide therefore adopts a kinked structure where Asp512 and Trp513 (−2 and −1 positions respectively) are directed upwards from the C-terminal Leu514 at position 0, and does not dock into the typical groove between helix αB and strand βB. Lastly, the structure of ZO1 PDZ domain 1 bound to a peptide from claudin-1 (KDYV) displays both conventional and unconventional binding within the same crystal lattice (Nomme et al., 2015). In each structure the Asp at −2 (Asp118) forms a hydrogen bond to the αB histidine His88, somewhat similarly to the way that Ser and Thr sidechains do this in typical Type I motifs. However, as for both the GRASP/mGluR1 (Elkins et al., 2010) and tamalin/tamalin (Sugi et al., 2007) complexes, the caveat is that these structures were determined using a short peptide fusion and there are adjacent crystal contacts that clearly influence the peptide conformations. Based on these comparisons, we propose that as a general rule Type III motifs with a Glu in the −2 position, as in SNX17 and myotilin, will be unable to bind PDLIM proteins in a conventional manner because of the steric interference between the longer Glu side chain and the conserved His sidechain in the PDZ domain αB helix (His63 in PDLIM7). Furthermore, the strictly conserved Arg sidechain in the βB strand of all PDLIM family PDZ domains (Arg17 in PDLIM7) (**Fig. 3G**) makes them particularly suited to specific binding of Type III motifs with acidic residues at both the −1 and −2 positions through complementary electrostatic interactions.

### Assessment of SNX17 interaction with PDLIM7 by mutagenesis

To confirm the mechanism of interaction between SNX17 and the PDZ domain of PDLIM7 a series of mutants were generated for both the PDZ domain and the SNX17 peptide with binding thermodynamics subsequently analysed by ITC (**Fig. 5; Table 1**). As expected, binding was abolished by the SNX17 mutation L470G at position 0 of the PDZbm (**Fig. 5A**). Similarly, the triple mutation of acidic side chains in SNX17 (D467S/E468S/D469S) also abolished binding, consistent with the important role of electrostatic interactions with PDLIM7, with Asp469 forming a stable contact with Arg17. In contrast, the single mutation E468R at the −2 position in SNX17 had only a modest effect on the interaction. A reciprocal mutation in the strictly conserved PDLIM7 R17D also prevented SNX17 binding, confirming the importance of the electrostatic contact between the Arg17 side chain and SNX17 Asp469 (**Fig. 5B and 5C**).

**Figure 5.**
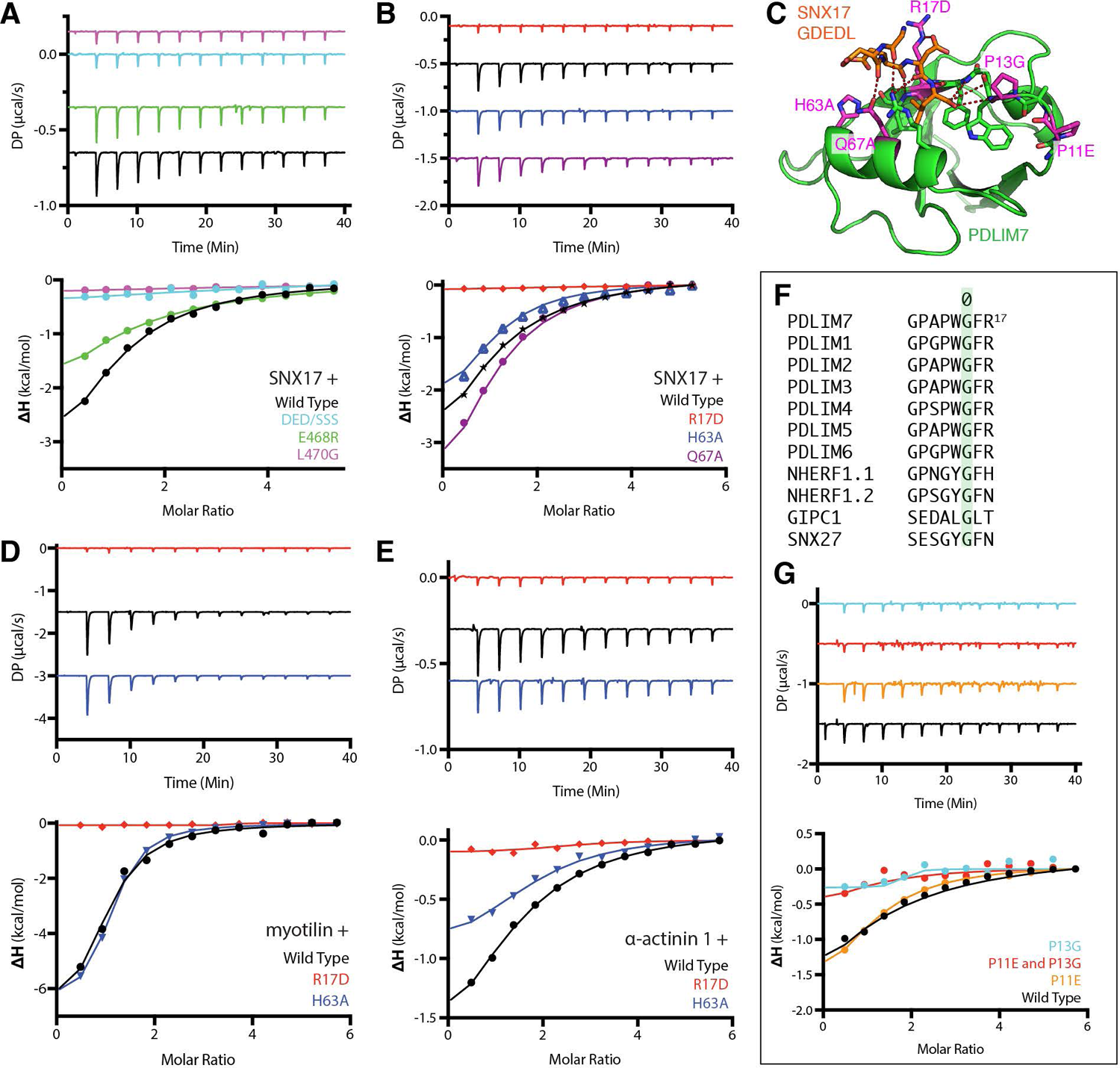
Mutagenesis and *in vitro* binding analysis confirms the binding mechanism for SNX17 interaction with the PDLIM7 PDZ domain. **(A)** Binding of the PDZ domain of PDLIM7 to mutants of the SNX17 peptide measured by ITC. The top panels show raw data while the bottom shows the normalised and integrated binding curves. **(B)** Binding of the wild-type SNX17 peptide to specific PDLIM7 mutants measured by ITC. (**C**) The sites of SNX17 point mutations are indicated in magenta on the structure of the PDLIM7-SNX17 complex. (**D**) Binding of the myotilin Type III PDZbm peptide to specific PDLIM7 mutants measured by ITC. (**E**) Binding of the α-actinin-1 Type I PDZbm peptide to specific PDLIM7 mutants measured by ITC. (**F**) Sequence alignment of the core binding loop preceding the βB strand of PDLIM family proteins and comparison to selected PDZ domains. A strictly conserved Gly residue is found in all PDZ domains (numbered ‘0’ for reference), but PDLIM proteins possess unique sequences upstream and downstream of this central residue, including an Arg at position +2 that coordinates with Acidic sidechains in the SNX17 peptide (see Fig. 3D), as well as Pro and Trp residues at the −2 and −1 positions respectively. **(G)** Binding of the wild-type SNX17 peptide to specific mutants of the PDLIM7 core loop sequence measured by ITC. P13G mutation at position −2 abolishes SNX17 binding, while the W14Y mutation at position −1 reduces binding affinity.

We next examined the importance of PDLIM7 His63 and Gln67 side chains in SNX17 binding. The structure indicates that His63, which is a conserved feature of nearly all PDZ domains and is important for conventional binding of Type I PDZbms, is not required for the unconventional binding of the SNX17 Type III PDZbm (**Fig. 3D**). Supporting this the H63A mutation had no effect on the SNX17 interaction (**Fig. 5B and 5C**). PDLIM7 Gln67 forms a hydrogen bond interaction with the backbone carbonyl of SNX17 Asp467 in the structure (**Fig. 3D**) but does not make a large contribution to the binding groove as seen for conventional PDZbm interactions (**Fig. 4**). Again, mutation of this residue (Q67A) has little effect on SNX17 binding (**Fig. 5B and 5C**). These results are all consistent with the unconventional SNX17 binding conformation. They further suggest that Type III motifs (especially those with a glutamate at the −2 position) will have a particular preference for unconventional binding of the PDLIM PDZ domains. Supporting this, we found that like SNX17 the binding of the known Type III ligand from myotilin (Elkins et al., 2010) is blocked by the R17D mutation in PDLIM7 but is unaffected by the H63A mutation (**Fig. 5D; Table 1**). In contrast the affinity of the known Type I motif from α-actinin-1 is perturbed by both mutations as expected for a canonical peptide interaction (**Fig. 5E; Table 1**).

The loop preceding the βB strand in PDLIM7 forms the core binding pocket for SNX17 and is composed of a sequence that is unique and highly conserved within the PDLIM protein family (**Fig. 5C and 5F**). In virtually all PDZ domains there is a conserved Gly residue (Gly15 in PDLIM7) within this binding loop, that provides main-chain amides for binding the C-terminal carboxylate-group of the incoming peptide, and this Gly residue is bookended by two hydrophobic side chains as part of a core consensus sequence x<ΦG<Φ (with the Gly residue numbered as ‘0’ for reference). (Lee and Zheng, 2010; Ye and Zhang, 2013). Uniquely, the PDLIM family invariably possess Pro, Trp and Phe side chains at the −2, −1 and +1 positions of this sequence respectively. While mutation of the Pro at position −4 in PDLIM7 (P11E) had little effect on SNX17 interaction, mutation of the Pro at position −2 (P13G) or double mutation of Pro at position −2 and −4 (P11E/P13G) mostly abolished binding to the SNX17 PDZbm, confirming the importance of Pro13 for the interaction (**Fig. 5G; Table 1**). Overall, our data indicates that the conserved sequence features of the PDLIM family provide a preferential platform for binding the SNX17 C-terminal sequence compared to other PDZ domain-containing proteins.

## Discussion

Here we have identified a novel association between SNX17, an adaptor for Commander-dependent endosomal trafficking, and the PDLIM family of actin-associated scaffold proteins. The interaction is mediated by the Type III PDZbm of SNX17 and appears to be highly specific for the PDZ domain of PDLIM proteins. Using NMR to determine the structure of the PDLIM7-SNX17 complex we show that their binding is dependent on unique sequence and structural properties of the interacting proteins, and this results in the SNX17 motif adopting an unconventional upright binding mode as opposed to occupying the conventional binding groove between the PDZ domain αB helix and βB strand seen for the vast majority of PDZ structures.

Our comparisons with other PDLIM PDZ domain complexes show that the unconventional PDLIM7-SNX17 structure shares the greatest similarity to the previous structure of PDLIM6 bound to the Type III motif of myotilin (PDB 4YDP, *unpublished*). Myotilin was first found to bind PDLIM7/Cypher via its C-terminal motif by yeast two-hybrid assay, and to colocalise with PDLIM6 at the Z-disc of muscle fibres (von Nandelstadh et al., 2009). It shares homology with myopalladin and palladin proteins that have related roles in in the Z-disc and other actin-dependent structures. These also have similar C-terminal Type III motifs that can bind to PDLIM family proteins (Hasegawa et al., 2010), as do additional Z-disc associated proteins of the myozenin family (von Nandelstadh et al., 2009). Several Type I motifs, such as the C-terminus of α-actinin-1, can adopt both conventional and unconventional binding modes when interacting with PDLIM family proteins (Elkins et al., 2010), and for PDZ structures more broadly both conventional and unconventional binding modes have been observed for each of the three major PDZbm classes (**Fig. 4**). However, as discussed above the functional significance of unconventional interactions seen for Type I motifs, as well as the conventional interaction seen for a single Type III motif, are still unclear due to the potential influence of short fusion sequences on their conformations within their crystal lattices. As a general observation, we propose that Type I motifs are most likely to preferentially adopt the conventional binding mode, while Type III motifs, especially those with a Glu sidechain at position −2, will necessarily bind in an unconventional orientation.

Both SNX17 and the PDLIM proteins appear to share a similar evolutionary distribution, present in metazoans from humans to worms but without clear homologues in plants, amoeba, or fungi (**Fig. 6A**). Moreover, both the C-terminal Type III PDZ motif of SNX17 and the core residues of the PDLIM PDZ domains across these species are highly conserved (**Fig. 6B**). This suggests that the association between these proteins is evolutionarily preserved. Interestingly, although PDLIM proteins are absent, SNX17 is also found in choanoflagellates (the closest single cell organisms to the metazoan kingdom), along with all components of Retriever, CCC and WASH complexes indicating that this endosomal machinery emerged early in animal evolution (**Fig. 6A**). Our study began with the hypothesis that because the C-terminal mutation of L470G in SNX17 could prevent its interaction with Commander there may be intermediate protein(s) containing a PDZ domain that link them together. The discovery that PDLIM proteins have high specificity for the SNX17 C-terminus prompts us to speculate that they may provide such a link, and their structures certainly suggest a potential protein scaffolding function (**Fig. 6C**). While this is a tempting idea there are several caveats. Firstly, although not tested exhaustively, we have not yet been able to demonstrate any direct interactions between immobilised recombinant GST-tagged PDLIM proteins and Retriever or CCC subunits in cell lysates (not shown). Secondly, there have been several proteome-wide studies that have detected an association of the various Commander subunits with each other, none have reported any clear evidence of PDLIM interactions (e.g. (Huttlin et al., 2017; Li et al., 2015; Phillips-Krawczak et al., 2015; Singla et al., 2019; Starokadomskyy et al., 2013; Wan et al., 2015)). In our own immunoprecipitation experiments using Retriever (**Fig. 1B**) and proteomic studies using other Commander subunits as baits (not shown) we also do not detect PDLIM proteins; although neither do we detect SNX17, which suggests that the SNX17-Commander interaction is more readily captured using SNX17 as the bait for unknown technical reasons. Lastly, because of the redundancy in the PDLIM family interactions with SNX17, it is difficult to test their potential role in linking SNX17 with Commander by simple depletion of the proteins by either siRNA or genetic knockout. Considering our proteomic and structural results, and with these caveats in mind, there are two distinct possibilities. Firstly, it is plausible that the PDLIM proteins do provide a link between SNX17 and Commander and their interaction with Commander simply remains to be characterised. Alternatively, while the C-terminal peptide of SNX17 binds to PDLIM proteins through their PDZ domain, it may also be binding independently to Commander through some other unknown mechanism. Further studies will be required to determine whether the PDLIM proteins play a direct role in regulating the interaction between SNX17 and Commander.

**Figure 6.**
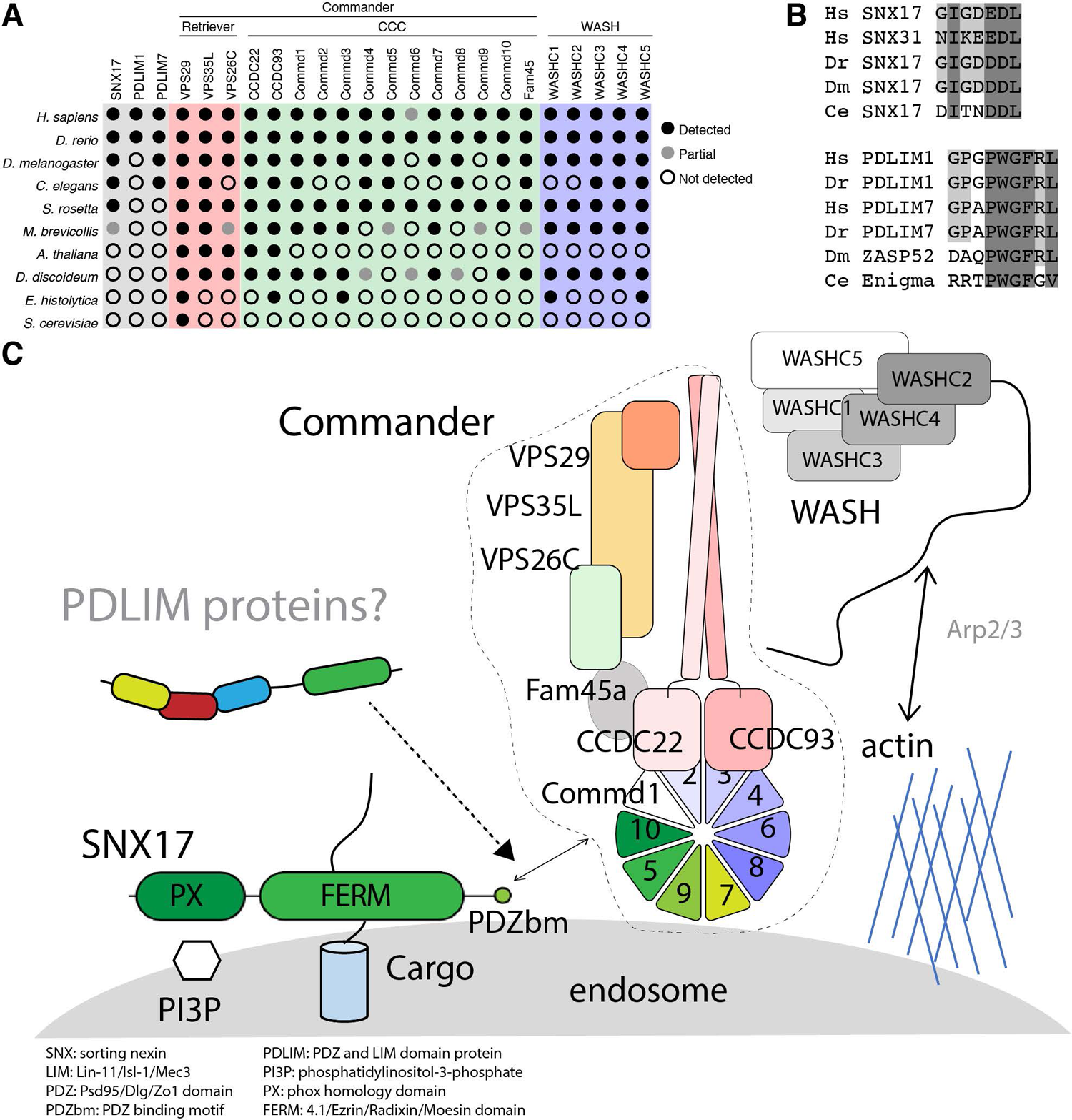
The evolutionary and functional relationship between PDLIM proteins, SNX17, Commander and the WASH complex. **(A)** Evolutionary conservation of SNX17, PDLIM proteins, the Commander complex and the WASH complex. Each protein was used to perform a BLAST search of the selected genomes ranging from metazoa (*Homo sapiens*, *Danio rerio*, *Drosophila melanogaster*, *Caenorhabditis elegans*), plants (*Arabidopsis thaliana*), simple eukaryotes (*Dictyostelium discoideum*, *Entamoeba histolytica*, *Saccharomyces cervisiae*) and the choanoflagellates as the nearest single celled animal precursors (*Salpingoaca rosetta*, *Monosiga brevicollis*). Only PDLIM1 (as a representative of the ALP subfamily) and PDLIM7 (as a representative of the Enigma subfamily) were used in these analyses. Black and grey circles indicate that full or partial homologues were identified. Closed circles indicate that no homologues were detected. (**B**) Sequence alignment across different species of the SNX17 C-terminus (top) and PDLIM loop region that binds to the C-terminus of the PDZbm. **(C)** Cartoon depicting the key interactions between PDLIM proteins, SNX17, Commander, and WASH proposed to be important for endosomal cargo sorting. Cargos with ΦxNxx[YF] motifs in their cytoplasmic tails are recruited via binding to the SNX17 FERM domain. We speculate that the PDLIM proteins may couple the C-terminal SNX17 PDZbm to the Commander assembly or otherwise modulate the assembly of actin-rich domains at the endosome required for SNX17-Commander mediated trafficking.

While SNX17, and the Retriever and CCC sub-complexes have established roles in endosomal trafficking, the PDLIM proteins have primarily been shown to be involved in the regulation of actin-mediated processes such as the formation of stress fibres and the Z-discs of muscle cells (Bauer et al., 2000; Katzemich et al., 2013; Klaavuniemi et al., 2009; Klaavuniemi et al., 2004; Liu et al., 2015; Tamura et al., 2007; te Velthuis and Bagowski, 2007; Vallenius et al., 2000; Vallenius et al., 2004; von Nandelstadh et al., 2009; Zheng et al., 2010). The PDLIM proteins have been implicated in endosomal regulation in one previous study, with the identification of an interaction between PDLIM1 and the glutamate receptor GluA1 that promotes their mutual colocalization at early endosomes, and enhances recruitment of actin to the endosomal compartment (Schulz et al., 2004). The transport of cargo proteins by SNX17 and Commander is highly dependent on localised actin dynamics controlled by the Arp2/3-activating WASH complex. The WASH complex is involved in endosomal trafficking and recruitment of both Commander and actin to the endosomal membrane (Bartuzi et al., 2016; Duleh and Welch, 2010; Gomez et al., 2012; Phillips-Krawczak et al., 2015; Simonetti and Cullen, 2019), and in turn the depletion of Commander negatively regulates this dynamic remodelling of actin, leading to the accumulation of filamentous actin on the endosome (Campion et al., 2018; Simonetti and Cullen, 2019; Singla et al., 2019). Our data therefore suggests the potential for interplay between PDLIM proteins, SNX17, Commander, WASH, and actin in regulating the formation of endosomal recycling domains that warrants further studies.

## Materials and methods

### Antibodies and peptides

All SNX17, α-actinin-1 and myotilin derived peptides were obtained from Genscript (USA). For immunofluorescence imaging we used goat anti-human VPS35 (Abcam; 10099), mouse anti-human EEA1 (BD Biosciences; 610457), Rabbit anti-human PDLIM7 (Novus; NBP1-84841) and TRITC-Phalloidin (Sigma; P-1951). Secondary antibodies used for immunofluorescence were (all from Invitrogen) AlexaFluor 488 donkey anti-rabbit IgG (A21206), AlexaFluor 568 anti-mouse IgG (A10037), AlexaFluor 568 anti-goat IgG (A11057), AlexaFluor 568 anti-rabbit IgG (A10042), AlexaFluor 647 anti-goat IgG (A21447). For Western blotting we used rabbit anti-human PDLIM7 (Proteintech; 10221-1-AP), rabbit anti-human VPS35L (Abcam; ab97889), rabbit anti-human CCDC93 (Proteintech; 16636-1-AP) and rabbit anti-human CCDC22 (Proteintech; 20861-1-AP). Secondary antibodies for Western blot analysis were AlexaFluor 680 Donkey anti-mouse IgG (Invitrogen; A10038), AlexaFluor 680 Donkey anti-rabbit IgG (Invitrogen; A11043) and AlexaFluor 800 donkey anti-rabbit IgG (Invitrogen; W10824).

### Protein enrichment for mass spectrometry

Biotinylated C-terminal SNX17 wild-type (^457^HGNFAFEGIGDEDL^470^) and SNX17 L470G (^457^HGNFAFEGIGDEDG^470^) peptides were resuspended at a concentration of 1 mg/mL in 400 μM ammonium hydroxide in a volume of 50 μL and incubated with 20 μL of streptavidin resin (Pierce Thermo) pre-washed with 100 μL PBS and equilibrated with 300 μL of Lysis Buffer. A total of 9 million of (or 10cm at 80% confluency) HEK293T cells were lysed in 1 mL of Lysis Buffer (1% (v/v) digitonin, 20 mM Tris (pH 7.4), 50 mM NaCl, 10% (v/v) Glycerol, 0.1 mM EDTA) and protein concentration measured by bicinchoninic acid assay (BCA; Pierce). The pre-washed peptides bound to streptavidin resin were incubated with lysate containing a total of 1 mg protein for 2 h at 4°C with rotating. Lysate was removed by centrifugation and the resin washed with 20x 300 μL of Wash Buffer (0.1% (v/v digitonin, 20mM Tris (pH 7.4), 60mM NaCl, 10% (v/v) Glycerol, 0.5 mM EDTA). Protein was eluted from resin using 30 μL of 6 M urea in 50 mM ammonium bicarbonate. Samples were reduced and alkylated with 50 mM tris(2-carboxyethyl)phosphine (TCEP) (ThermoFisher) and 500 mM chloroacetamide (Merck) for 30 min at 37°C with shaking. Eluates were diluted to 2 M urea in 50 mM ammonium bicarbonate and digested with 1 μg of trypsin (ThermoFisher) overnight at 37°C. The digested material was acidified to a final concentration of 1% (v/v) trifluoracetic acid (TFA) and peptides were desalted with in-house made stage-tips containing 2x plugs of 3M™ Empore™ SDB-XC substrate (SUPELCO) (Hock et al., 2020; Kulak et al., 2014). Stage-tips were activated with 100 % acetonitrile (ACN) and washed with 0.1% (v/v) TFA, 2% (v/v) ACN prior to binding the peptides. Samples were eluted in 80% ACN, 0.1% TFA and dried completely in a SpeedVac. Peptides were reconstituted in 0.1% TFA, 2% ACN, and transferred to autosampler vials for analysis by high-performance liquid chromatograph (HPLC)-coupled tandem mass spectrometry (MS/MS) on an LTQ Orbitrap Elite (ThermoFisher) connected to an Ultimate 3000 HPLC (Dionex) and nano-electrospray ionisation (ESI) interface by the Melbourne Mass Spectrometry and Proteomics Facility (MMSPF). Instrument parameters were as follows. Peptides were loaded onto a trap column (Dionex-C18 trap column 75 μm × 2 cm, 3 μm, particle size, 100 Å pore size; ThermoFisher Scientific) at 5 μl/min before switching the pre-column in line with the analytical column (Dionex-C18 analytical column 75 μm × 50 cm, 2 μm particle size, 100 Å pore size; ThermoFisher Scientific). The separation of peptides was performed at 300 nl/min using a 65 min (total run length) non-linear ACN gradient of buffer A (0.1% formic acid, 2% ACN, 5% DMSO) and buffer B (0.1% formic acid in ACN, 5% DMSO). Data was collected in Data Dependent Acquisition (DDA) mode using m/z 300–1650 as MS scan range, rCID for MS/MS of the 20 most intense ions. Dynamic exclusion with of 30 seconds was applied for repeated precursors.

### Mass spectrometry and data analysis

Raw files were analysed using the MaxQuant platform (Cox and Mann, 2008), version 1.6.10.43 against canonical, reviewed and isoform variants of human protein sequences in FASTA format (Uniprot, January 2019). The default settings: “LFQ” and “Match between runs” were enabled. N-terminal acetylation and methionine oxidation were set as variable modifications while cysteine carbamidomethylation was specified as a fixed modification. Computation of SNX17_DEDL_/SNX17_DEDG_ enrichment was performed in Perseus (version 1.6.10.43) (Tyanova et al., 2016). Peptides labelled by MaxQuant as ‘only identified by site’, ‘reverse’ or ‘potential contaminant’ were removed and only those proteins quantified based on >1 unique peptide were considered for further analysis. LFQ intensities were log2 transformed and rows having less than 3 valid values in the enrichment group (SNX17_DEDL_) were removed and the missing values in the SNX17_DEDG_ control group were imputed to values consistent with the limit of detection. The mean log_2_ LFQ intensities for proteins detected in each experimental group, along with p-values, were calculated using a two-sided two-tailed t-test. Significance was determined by permutation-based FDR statistics (Tyanova et al., 2016) where the s0 factor was iteratively modified to exclude all identifications enriched in the control (SNX17_DEDG_) experiment, yielding an s0 of 1 at 1% FDR.

### Molecular biology and cloning

The PDZ domains of human PDLIM1 (residues 1-84), PDLIM2 (1-84), PDLIM3 (1-84), PDLIM4 (1-84), PDLIM5 (1-84), GIPC1 (133-213), PTPN3 (510-582), NHERF1.1 (14-94), NHERF1.2 (154-234), and SNX27 (43-136), full-length sequence of VPS26C plus the PDLIM7 PDZ (1-84) fusion with SNX17 (457-470) peptide were artificially synthesised as codon optimised constructs and cloned into pGEX6P-1 between BamHI and XhoI by Gene Universal for bacterial expression with an N-terminal GST-tag and a Prescission cleavage site. The PDLIM7-SNX17 fusion was designed to include the SNX17 residues 457-470 with a preceding flexible linker (GASAGASAGASA^457^HGNFAFEGIGDEDL^470^). The gene sequences for the PDZ domain of human PDLIM7 (1-84) and various point mutants were synthesised as codon optimised constructs and cloned into pET30b(+) by Genscript (USA) for bacterial expression with an N-terminal 6His-tag.

For mammalian expression, pEGFP-C1-SNX17 WT and mutants, pXLG3-GFP-SNX17 and pEGFP-C1-VPS26C vectors were made previously (McNally et al, 2017). To express PDLIM7-GFP, cells were transfected with pcDNA-PDEST47 PDLIM7-GFP (Elbediwy et al., 2018). PDLIM1 and PDLIM7 were both also cloned into a pEGFP-N1 vector between HindIII and BamHI for mammalian expression with C-terminal GFP tags. For lentiviral transduction of RPE-1 cells, human CCDC22 was first cloned into pEGFP-C1 (Clontech) between Sal1 and BamH1. GFP-CCDC22 was then excised using Nhe1 and BamH1 and ligated into pXLG3 digested with Spe1 and BamH1.

### Recombinant protein expression and purification

Bacterial expression vectors were transformed into *Escherichia coli* BL21 (DE3) competent cells (New England Biolabs). The bacterial cultures were grown in LB until they reached OD_600nm_ 0.8. Cultures were immediately induced with 0.8 mM ispropylthio-β-galactoside (IPTG) before temperature was reduced to 21°C and cultures were allowed to grow for 18 h. The cells were harvested by centrifugation at 6000x g for 5 min and cultures were resuspended in lysis buffer (500 mM NaCl, 20 mM Tris (pH 7.4), 2 mM β-mercaptoethanol, 50 mg/mL benzamidine, and 100 units DNAseI). Cells expressing 6His-tagged proteins were supplemented with 5 mM Imidazole (pH 8.0) before cell lysis. Cells were lysed by mechanical disruption at 35 kpsi using a Constant systems cell disruptor before lysate clarification by centrifugation (50,000x g for 45 min at 4°C). Proteins were then purified using affinity chromatography from the clarified lysate.

His-tagged proteins were purified using TALON resin (Bio-strategy Pty Limited) in a gravity flow column and eluted in a buffer containing 500 mM imidazole, 200 mM NaCl, 20 mM Tris (pH 7.4), and 2 mM β-mercaptoethanol. GST-tagged proteins were purified using glutathione-Sepharose (GE health) gravity flow column and eluted using 10 mM glutathione, 200 mM NaCl, 20 mM Tris (pH 7.4) and 2 mM β-mercaptoethanol. In instances where a cleaved protein was required 0.5 mg of purified Prescission protease was added to the buffer above and allowed to incubate overnight before protein elution. Finally, proteins were subjected to size exclusion chromatography using either a Superdex 200 10/300 or Superdex 75 10/300 column attached to an AKTA Pure system (GE Healthcare).

In order to produce the isotopically labelled protein a modified version of the Marley method was used (Marley et al., 2001). In brief, overnight culture was transferred into 1 L flasks and grown in normal LB media till OD_600_ reached 0.8. Cells were then harvested using sterile centrifugation tubes at 6000x g for 5 min and resuspended in 1 L M9-A salts (110 mM KH_2_PO_4_, 260 mM Na_2_HPO_4_ and 42.7 mM NaCl). Cells were spun down again at 6000x g for 5 mins and resuspended in 250 mL of M9-B (110 mM KH_2_PO_4_, 260 mM Na_2_HPO_4_, 42.7 mM NaCl, 1 x MEM Vitamin solution (Sigma), 2 mM MgSO_4_ 1 μM CaCl_2_, 18.7 mM ^15^NH_4_Cl, 22.2 mM ^13^C-D-glucose). Resuspended cells were induced with 1 mM IPTG and grown over night at 18°C. Purification was conducted as above, and the labelled protein was gel filtered into a buffer containing: 75 mM NaCl bis-tris (pH 6.0) and 4 mM dithiothreitol and 5% v/v D_2_O.

### Isothermal titration calorimetry (ITC)

The affinities of PDZ domain and VPS26C interactions with SNX17, α-actinin-1 and myotilin peptides were determined using a Microcal PEAQ instrument (Malvern, UK). Experiments were performed in 50 mM Tris-HCl (pH 7.4), 100 mM NaCl. The native and mutant peptides at 700 μM were titrated into 25 μM of PDZ domains in 13 x 3.22 μL aliquots at 37°C. In the case of the DSCR3, PDLIM7_LIM interaction x mM of DSCR3 was titrated into 20 mM PDLIM7_LIM at 25°C. The dissociation constants (*K*_d_), enthalpy of binding (ι1*H*) and stoichiometries (N) were obtained after fitting the integrated and normalized data to a single-site binding model. The apparent binding free energy (ι1*G*) and entropy (ι1*S*) were calculated from the relationships ι1*G* = RTln(*K*_d_) and ι1*G* = ι1*H* - Tι1*S*. All experiments were performed at least in triplicate to check for reproducibility of the data.

### Nuclear magnetic resonance (NMR) spectra collection

NMR spectra of the PDLIM7 PDZ domain fused to the SNX17 peptide were acquired in 75 mM NaCl, 20 mM bis-tris (pH 5.8), 4 mM DTT and 5% D_2_O at a protein concentration 1 mM using a Shigemi NMR tube (Sigma Aldrich). Data was collected on an ultrahigh field Bruker 900 MHz NMR spectrometer equipped with a triple resonance cryogenic probe. Backbone resonance assignments were obtained from a 2D ^1^H-^15^N-HSQC, 2D ^1^H-^13^C-HSQC, 3D HNCACB, 3D CBCA(CO)NH, 3D HNCO, and 3D HBHA(CO)NH spectra, whereas side chains were determined using 3D HccoNH and 3D hCcoNH.

### NMR spectral analysis and structural calculations

Structure constraints were calculated from ^13^C (separate spectra for aliphatic and aromatic regions) and ^15^N-NOESY HSQC spectra. Peak data was calculated using the CCPNMR software (Skinner et al., 2016) and the final structural calculation was made by CYANA 3.98-5 (Guntert and Buchner, 2015), using backbone dihedral restraints calculated from TALOS-N (Shen and Bax, 2015). The parameters used in CYANA generated 200 structures of which the 20 lowest energy models were selected. The final structure statistics can be found in **Table 2**.

### Cell culture

HEK293, HEK293T, HeLa and RPE1 cells were maintained in DMEM (D5796; Sigma-Aldrich) plus 10% fetal calf serum (F7524; Sigma-Aldrich) under standard conditions. These cell lines were obtained from America Type Culture Collection (ATCC). Parental and stable cells lines were negative for mycoplasma by DAPI staining. Lentivirus particles for producing stably expressing cell lines were generated in HEK293 cells using the pXLG3 vector to carry the GFP tagged CCDC22. Cells were transfected with DNA using polyethylenimine (Sigma-Aldrich). Virus was harvested from the growth media 72 hr post transfection.

For stable transduction with lentivirus, RPE-1 cells were seeded at 50,000 per well in six well plates. The cells were then incubated under normal conditions with titrations of viral supernatant for 72 hr. Cells were then passaged and expression of the GFP tagged protein of interest assessed by Western blot analysis. Cell lines that displayed expression levels close to endogenous levels of protein were selected. GFP-SNX17 stable RPE-1 cells used in this study were made previously (McNally et al., 2017).

### GFP Traps

PEI (polyethylenimine) was used to transfect HEK 293 cells with constructs for GFP traps. An aqueous 10µg/ml stock of linear 25kDa PEI (Polysciences, catalogue number 23966-2) was used for the transfection. For 10 cm or 15 cm, 2.5ml or 5ml Opti-MEM^®^ was added to 2 separate sterile tubes respectively. In the first Opti-MEM^®^ containing tube, 10µg or 15µg DNA was added for 10cm or 15cm dishes respectively. To the second tube, 3µl or 4.5µl (for 10cm and 15cm dishes respectively) of PEI was added and the contents of the tube mixed by vortexing. The Opti-MEM^®^/PEI mixture was then filter sterilised by filtering through a 0.2µm filter. Sterilised PEI/Opti-MEM^®^ was then added to the Opti-MEM^®^ /DNA mixture and the tube was mixed by vortexing. The mixture was left to incubate at room temperature for 20 minutes. HEK 293 cells were washed in PBS and then the transfection mixtures were carefully added to the cell dishes. HEK 293 cells were incubated with the transfection mixture, under normal growth conditions, for 4 hours. The transfection media was removed at the end of the incubation period and replaced with normal growth media. Cells were further incubated for another 48 hours prior to experimental use.

Dishes containing cells expressing GFP or GFP tagged proteins were placed on ice. The cell media was removed and the cells were washed three times with ice cold PBS (Sigma). Cells were lysed with lysis buffer (20 mM Hepes pH 7.2, 50 mM potassium acetate, 1 mM EDTA, 200 mM D-sorbitol, 0.1% Triton X-100, 1x protease cocktail inhibitor (Roche)). 500µl or 1 ml of lysis buffer was used per 10 cm or 15 cm dish respectively. Lysis was aided through the use of a cell scraper. The lysates were then cleared by centrifugation at 13,200 rpm for ten minutes at 4°C. 15 µl of GFP-trap beads (Chromotek; gta-20) were equilibrated in lysis buffer, through three rounds of pelleting beads and re-suspension in lysis buffer, prior to adding cleared cell lysate. 10% of cell lysate was retained for input analysis. GFP-trap beads and lysates were incubated on a rocker at 4°C for 1 hour. Following incubation, GFP-trap beads were pelleted by centrifugation at 4000 rpm, for 30 seconds at 4°C. Supernatant was then removed and beads were washed a further three times in lysis buffer through rounds of re-suspension and pelleting. After the final wash, all lysis buffer was removed from the GFP trap beads. Beads were then either stored at −20°C or processed for SDS-PAGE analysis.

### Immunofluorescence

RPE-1 or Hela cells grown on 13 mm coverslips were washed with PBS before being fixed in ice cold 4% formaldehyde in PBS for 25 min. Cells were permeabilised in 0.1% Triton X-100 (Sigma) for 6 min. The cells were then blocked with 1% (w/v) bovine serum albumin (BSA) in PBS for 15 min at room temperature. Primary antibodies were diluted in 1% (w/v) BSA in PBS and samples were incubated for 1 hr at room temperature. The samples were then washed in PBS and then incubated with Alexa Fluor conjugated secondary antibody and 0.2 mM DAPI for 30 min at room temperature. Coverslips were mounted in Mowiol-DABCO mounting medium (Sigma). Cells were visualised using a Leica TCS SP5 X confocal microscope (Leica Biosystems) (**Fig. 1D; Fig S1A, B, C**) or a Zeiss LSM880 fast Airyscan (ZEISS microscopy) (**Fig. S1D**).

## Supporting information

Table S1 Proteomic data

## Acknowledgements

This work was supported by funds from the National Health and Medical Research Council (NHMRC) to RG and DAS (GNT1156732). MDH is supported by a Postgraduate Research Award from the Australian Institute of Nuclear Science and Engineering (AINSE; ALNSTU12277). BMC and DAS are supported by an NHMRC Fellowships (GNT1136021 to BMC; GNT1140851 to DAS). MM was supported by the Australian Research Council (ARC grant DP190101177) and the NHMRC (GNT1162597). PJC received funding from the MRC (MR/P018807/1), Wellcome Trust (104568/Z/14/Z and 220260/Z/20/Z), Lister Institute of Preventive Medicine, and the Royal Society (RSRP/R1/211004). We would like to thank Barry Thompson for the kind gift of the pcDNA-PDEST47 PDLIM7-GFP plasmid. We thank the Bio21 Mass Spectrometry and Proteomics Facility for the provision of instrumentation. This study made use of the ACRF Cancer Biology Imaging Facilities at the IMB (University of Queensland).

## Author contributions

Conceptualization, MDH, RG, DAS, BMC.; Methodology, MDH, JS, KEM, PJC, MM, RG, DAS, BMC; Investigation, MDH, JS, KEM, CM, MM, RAG, BMC; Writing – Original Draft, MDH, BMC; Writing – Review & Editing, MDH, JS, KEM, PJC, MM, RG, DAS, BMC; Funding Acquisition, PJC, MM, RG, DAS, BMC; Supervision, PJC, MM, RG, DAS, BMC.

## Data Availability

Coordinates and NMR chemical shift data for the PDLIM7 domain in complex with SNX17 have been deposited respectively at the Protein Data Bank (PDB) with accession code 7RM8 and Biological Magnetic Resonance Bank (BMRB) with accession code 30937. The mass spectrometry proteomics data will be deposited in the ProteomeXchange Consortium via the PRIDE (Perez-Riverol et al., 2019) partner repository upon publication. All the relevant raw data related to this study is available from the corresponding authors on request.

## Supplementary Information

**Figure S1.**
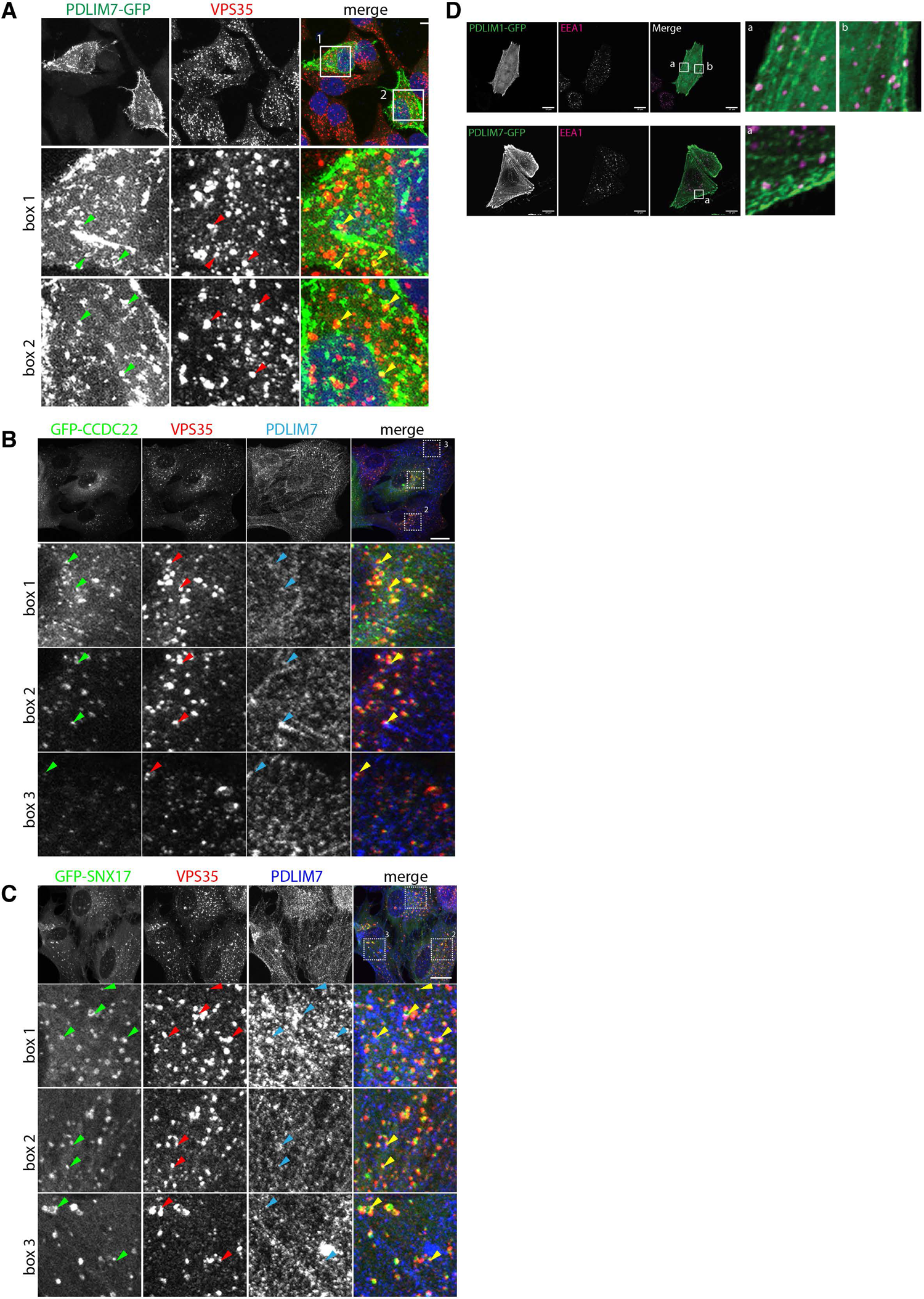
Localisation of PDLIM7 in cells. (**A**) Localisation of PDLIM7-GFP expressed in HeLa cells. (**B,C**) Localisation of endogenous PDLIM7 in lentiviral transduced RPE1 cells expressing GFP-CCDC22 or GFP-SNX17 respectively. Arrows indicate instances of PDLIM7 puncta overlapping with endosomal compartments. (**D**) Localisation of PDLIM1-GFP expressed in HeLa cells with PDLIM7-GFP shown for comparison. Scale bars represent 10µm.

**Figure S2.**
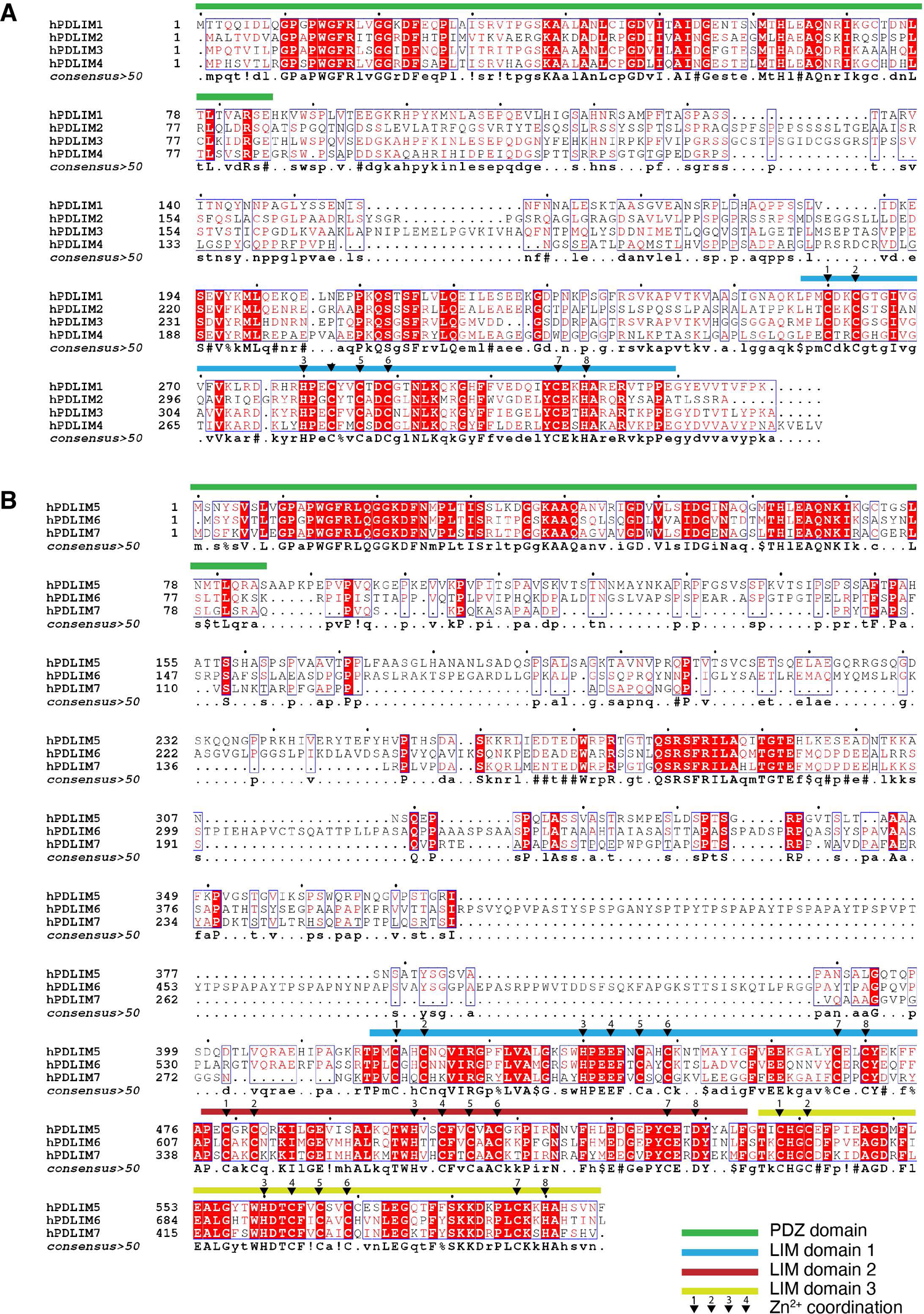
PDLIM family sequence alignments. Sequence alignments of the PDLIM family members from the **(A)** single LIM domain-containing ALP sub-family and the **(B)** triple LIM domain-containing Enigma sub-family. Each LIM domain contains eight residues indicated by numbered triangles that coordinate two Zn^2+^ ions. Sidechains 1-4 coordinate one Zn^2+^ ion and sidechains 5-8 coordinate a second Zn^2+^ ion.

**Figure S3.**
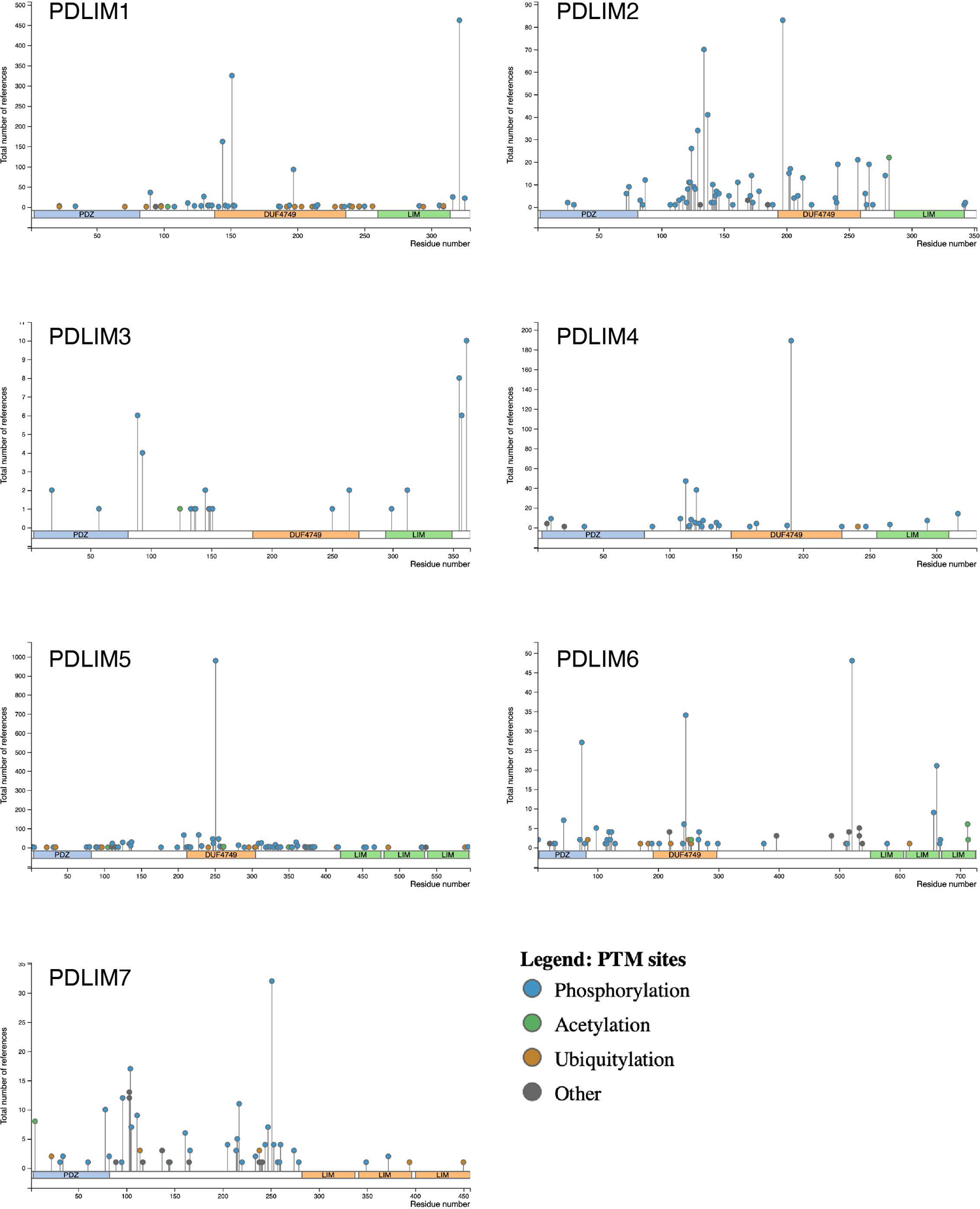
Post-translational modifications in the PDLIM family proteins. Sites of phosphorylation, acetylation and ubiquitination of PDLIM family members were identified using the PhosphositePlus database (https://www.phosphosite.org) (Hornbeck et al., 2015).

**Figure S4.**
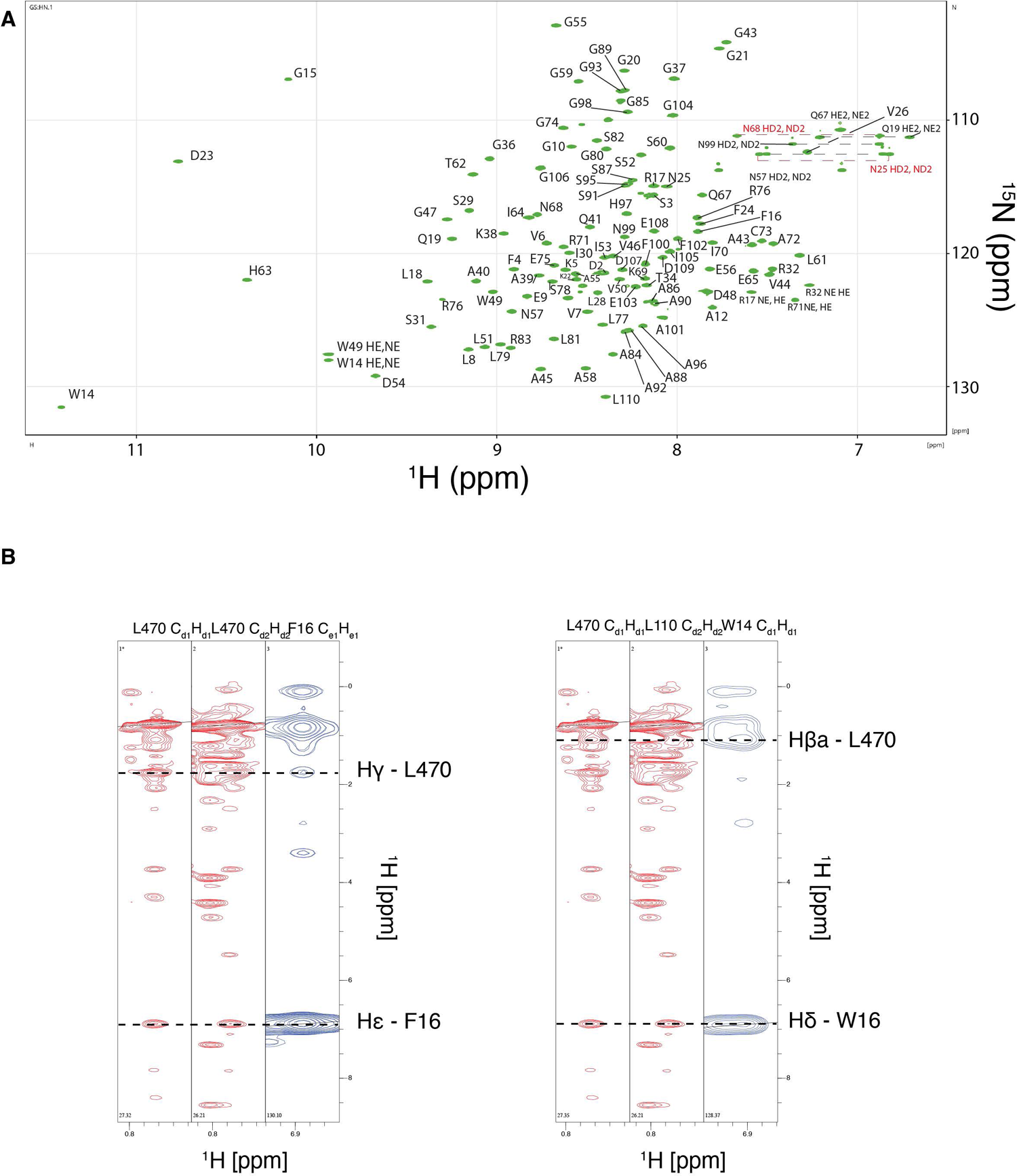
NMR spectra of the PDLIM7-SNX17 fusion protein. **(A)** ^15^N-^1^H HSQC spectra of ^15^N/^13^C-labeled PDLIM7-SNX17 fusion protein. **(B)** NOESY spectra of the PDLIM7-SNX17 fusion protein indicating contacts between Leu470 and the Trp14 and Phe16 aromatic sidechains.

---

**Table S1 Proteomic results of SNX17 C-terminal peptide interactions from HEK293 cells.**

## Key Resources Table

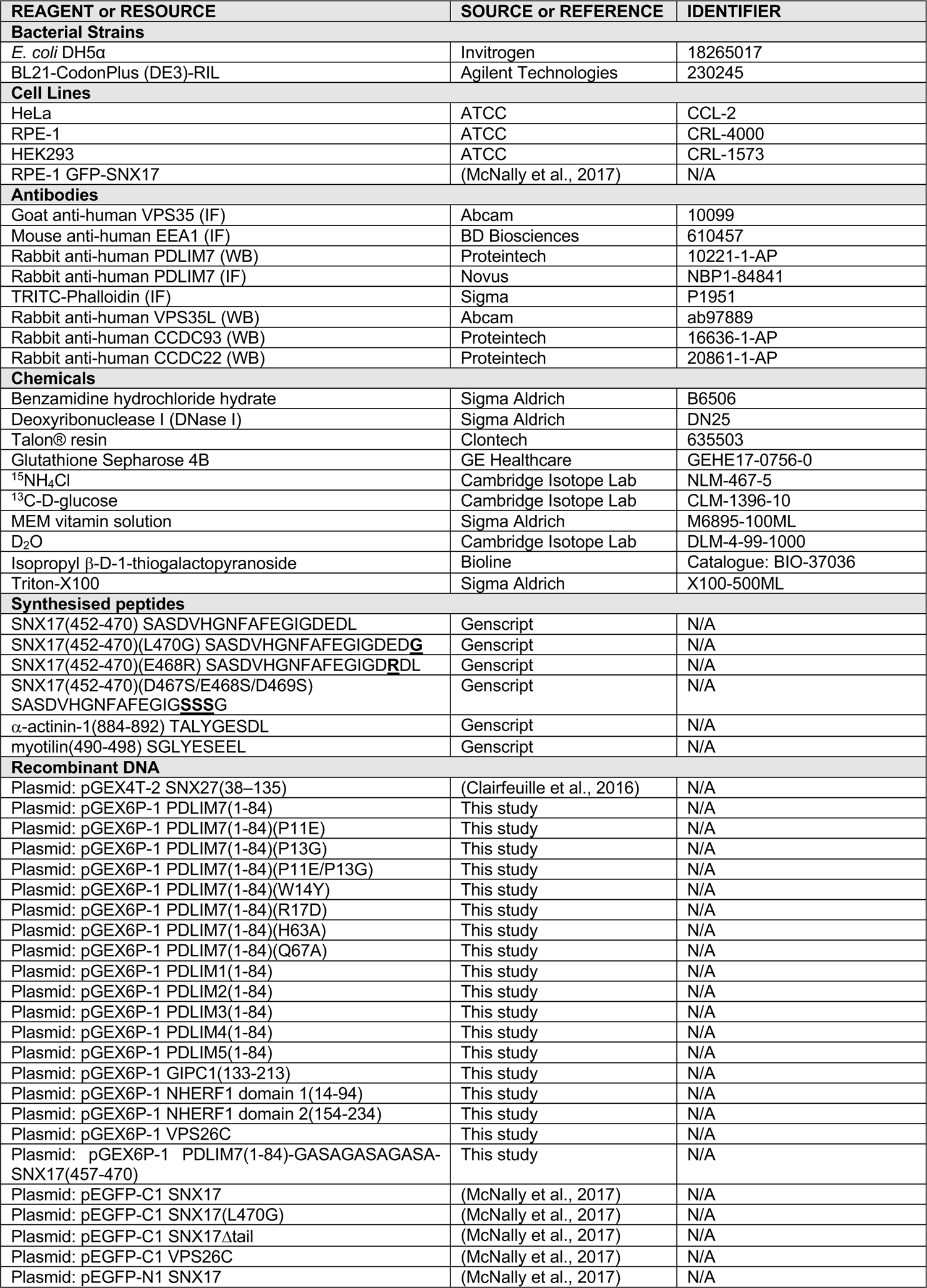
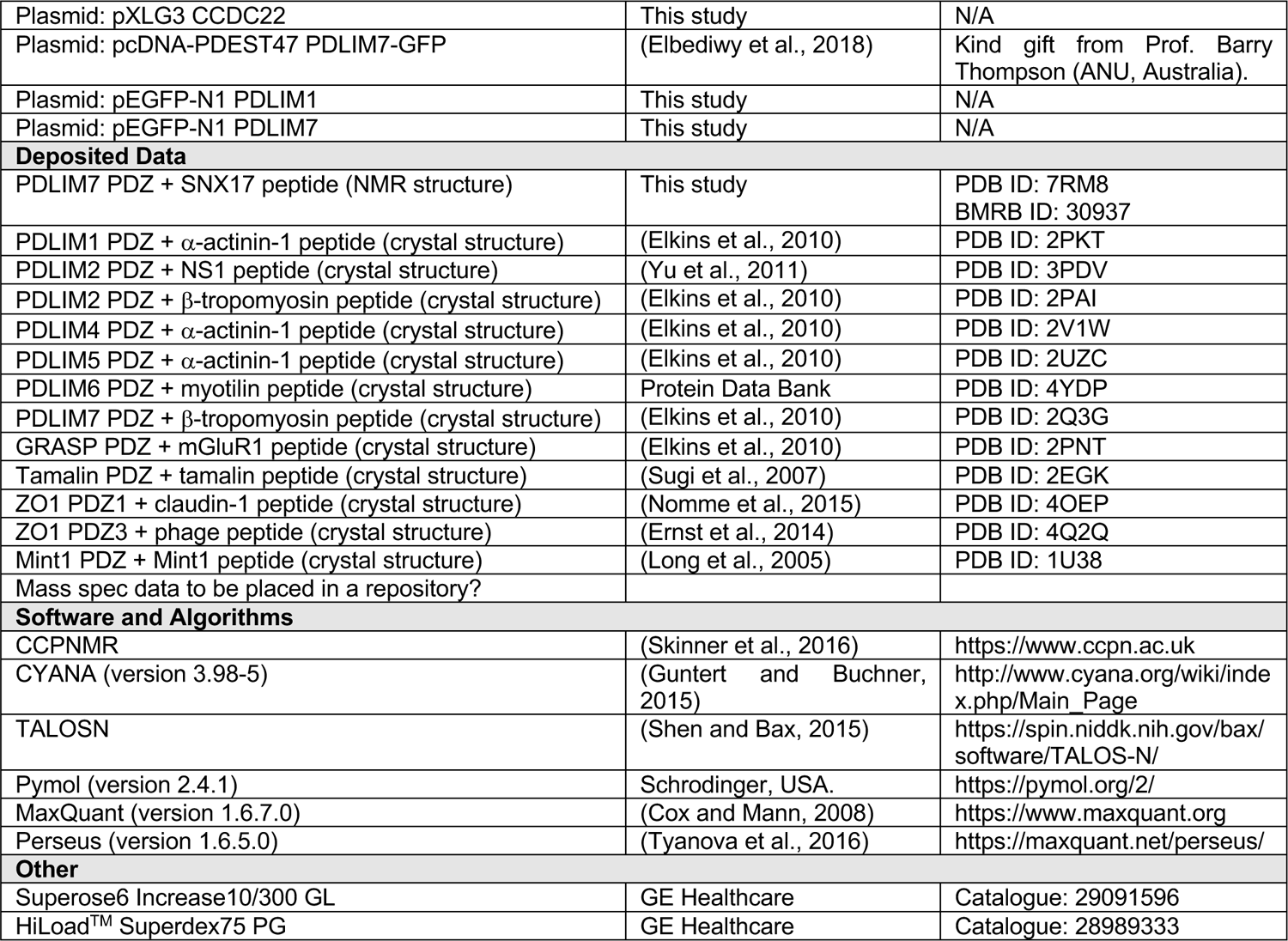

